# Genetic screen to saturate guard cell signaling network reveals a role of GDP-L-fucose metabolism in stomatal closure

**DOI:** 10.1101/2020.06.04.134353

**Authors:** Cezary Waszczak, Triin Vahisalu, Dmitry Yarmolinsky, Maija Sierla, Olena Zamora, Marina Leal Gavarrón, Julia Palorinne, Ross Carter, Ashutosh K. Pandey, Maris Nuhkat, Melanie Carmody, Tuomas Puukko, Nina Sipari, Airi Lamminmäki, Jörg Durner, Dieter Ernst, J. Barbro Winkler, Lars Paulin, Petri Auvinen, Andrew J. Fleming, Jarkko Salojärvi, Hannes Kollist, Jaakko Kangasjärvi

## Abstract

Guard cells regulate plant gas exchange by controlling the aperture of stomatal pores. The process of stomatal closure involves a multi-input signaling network that governs the activity of ion channels, which in turn regulate guard cell turgor pressure and volume. Here we describe a forward genetic screen to identify novel components involved in stomatal movements. Through an ozone-sensitivity approach combined with whole-rosette gas exchange analysis, 130 mutants of established stomatal regulators and 76 novel mutants impaired in stomatal closure were identified. One of the novel mutants was mapped to MURUS1 (MUR1), the first enzyme in *de novo* GDP-L-fucose biosynthesis. Defects in synthesis or import of GDP-L-Fuc into the Golgi apparatus resulted in impaired stomatal closure to multiple stimuli. Stomatal phenotypes observed in *mur1* were independent from the canonical guard cell signaling and instead could be related to altered mechanical properties of guard cell walls. Impaired fucosylation of xyloglucan, N-linked glycans and arabinogalactan proteins did not explain the aberrant function of *mur1* stomata, however our data suggest that the stomatal phenotypes observed in *mur1* can at least partially be attributed to defective dimerization of rhamnogalactouronan-II. In addition to providing the genetic framework for future studies on guard cell signaling, our work emphasizes the impact of fucose metabolism on stomatal movement.

## INTRODUCTION

Stomata are epidermal pores surrounded by pairs of guard cells that balance the loss of water and uptake of CO_2_ for photosynthesis. Guard cells respond to multiple environmental factors e.g. light, CO_2_ concentration, drought, low humidity, pathogens and air pollutants such as ozone (O_3_), to optimize transpiration or prevent the entry of the pathogens into the leaf tissue. Accumulation of reactive oxygen species (ROS) in the apoplast of guard cells and subsequent activation of plasma membrane Ca^2+^_in_ channels are among the first events associated with execution of stomatal closure (McAinsh et al., 1996; Pei et al., 2000; Kwak et al., 2003).

Depending on the stimulus, the apoplastic ROS are generated by NADPH oxidases (Kwak et al., 2003; Kadota et al., 2014), apoplastic peroxidases and amine oxidases (Sierla et al., 2016), however, external application of ROS alone is sufficient to initiate the process of stomatal closure (Price, 1990; McAinsh et al., 1996; Kollist et al., 2007). Hydrogen peroxide (H_2_O_2_) is the most stable form of ROS (Waszczak et al., 2018). Apoplastic perception of H_2_O_2_ involves activation of HYDROGEN PEROXIDE-INDUCED Ca^2+^ INCREASES1 (HPCA1) leucine-rich repeat receptor kinase which is necessary for activation of Ca^2+^_in_ channels (Wu et al., 2020). The subsequent rise in cytoplasmic Ca^2+^ concentration activates multiple Ca^2+^-dependent protein kinases (Maierhofer et al., 2014; Brandt et al., 2015) that together with OPEN STOMATA1 (OST1) kinase (Geiger et al., 2009; Lee et al., 2009) phosphorylate and activate guard cell anion channels SLOW ANION CHANNEL-ASSOCIATED1 (SLAC1), QUICK-ACTIVATING ANION CHANNEL1 (QUAC1) and SLAC1 HOMOLOGUE3 (SLAH3; for full list of kinases see Sierla et al., (2016)). The activation of guard cell anion channels, accompanied by deactivation of H^+^-ATPase1 (AHA1; Merlot et al., 2007) leads to membrane depolarization and activation of K^+^_out_ channels (Hedrich, 2012). The efflux of ions into the apoplast leads to a decrease of osmotic pressure inside the guard cells which provokes an efflux of H_2_O from the guard cell cytoplasm and vacuole. The consequent drop in guard cell turgor pressure results in closure of stomatal pores (Franks et al., 1998).

As evidenced by measurements of guard cell volume and turgor pressure performed in broad bean (*Vicia faba*), during stomatal opening guard cell turgor pressure rises from as low as 0.3 MPa to 5 MPa which is accompanied by a 30-40 % increase in guard cell volume (Franks et al., 2001). Importantly, the expansion and flexing of guard cells has to overcome the turgor pressure of the subsidiary cells (for a detailed discussion of the role of subsidiary cells see Lawson and Matthews, (2020)). On the other hand, stomatal closure involves a significant decrease in guard cell volume and surface area (Shope et al., 2003). To allow these volume and pressure changes, the guard cell walls must have a high degree of plasticity, which must be determined by wall structure. While differences between taxa exist (Popper, 2008), plant primary cell walls are typically composed of cellulose, hemicelluloses (xyloglucan, xylan, mannan), structural proteins, and pectins such as homogalactouronan (HG), rhamnogalactouronan-I (RG-I) and rhamnogalactouronan-II (RG-II) that determine the cell wall elasticity (Liepman et al., 2010). However, the relative content of these components varies between cell types and depends on the developmental stage. Moreover, the spatial distribution of different cell wall components is not uniform and reflects the mechanical needs of the respective cell type. Guard cells are an excellent example of such specialization. In comparison to epidermal cells, the guard cell walls are significantly thicker, devoid of highly methyl-esterified HG and rich in un-esterified HG (Amsbury et al., 2016; Merced and Renzaglia, 2018). The degree of methyl esterification is inversely correlated with the ability for Ca^2+^-mediated crosslinking, which leads to a more rigid cell wall, as well as susceptibility to degradation by polygalacturonases that leads to cell wall loosening (Levesque-Tremblay et al., 2015). Plants deficient in pectin demethylesterification exhibit defects in stomatal closure (Amsbury et al., 2016) which suggests that pectin crosslinking/degradation has a profound effect on the execution of this process. Further, targeted enzymatic digestion of arabinans that typically constitute the side chains of RG-I inhibits stomatal movements, and this effect can be counteracted by subsequent digestion or depolymerization of HG (Jones et al., 2003). Based on this observation, Jones et al., (2003) proposed a model in which the arabinan side chains of RG-I prevent crosslinking of HG strands which otherwise increases the cell wall rigidity, making it less capable to react to changes in guard cell turgor. Further, as observed already in the first half of the 20^th^ century (see Shtein et al., (2017) for recent visualization) the cellulose microfibrils within the guard cell walls fan out radially from the pore to provide a hoop reinforcement that limits the increase in guard cell radius and promotes guard cell elongation during the stomatal opening (Woolfenden et al., 2017). Moreover, as inferred from the atomic force microscopy (AFM)-based studies of guard cells, the stiffness of guard cell walls is not uniform; the most rigid areas are localized at the poles of guard cells and ventral walls directly surrounding the pore (Carter et al., 2017). During stomatal opening the polar fixing prevents the increase in the length of stomatal complex and forces the elongating guard cells to bend, leading to an increase in pore aperture (Carter et al., 2017). Taken together, it is clear, that the mechanics of the guard cell wall has a profound role in the execution of stomatal movements (Rui et al., 2018; Woolfenden et al., 2018).

The synthesis of cell wall glycan polymers relies on the availability of nucleotide sugars that constitute the activated precursor forms serving as a donor of sugar moieties (Bar-Peled and O’Neill, 2011). The importance of nucleotide sugar synthesis and transport is exemplified by the requirement of GDP-L-fucose for proper growth and development (Reiter et al., 1993; Reiter et al., 1997; O’Neill et al., 2001; Van Hengel and Roberts, 2002; Rautengarten et al., 2016), resistance to pathogens (Zhang et al., 2019) and freezing tolerance (Panter et al., 2019). The synthesis of GDP-L-Fuc is initiated by GDP-D-mannose 4,6-dehydratases (GMD1) and MURUS1 (MUR1/GMD2) that catalyze the conversion of GDP-D-mannose to GDP-4-keto-6-deoxy-D-mannose (Figure 1; Bonin et al., 1997; Bonin et al., 2003). In leaf tissues, MUR1 is the major GMD isoform and the level of L-Fuc observed in cell walls of *mur1* mutants is reduced by approximately 98% as compared to wild type plants (Reiter et al., 1993). In aerial organs GMD1 is expressed only in stipules and pollen grains (Bonin et al., 2003) and plays a minimal role in the synthesis of GDP-L-Fuc in leaf tissue.

**Figure 1.**
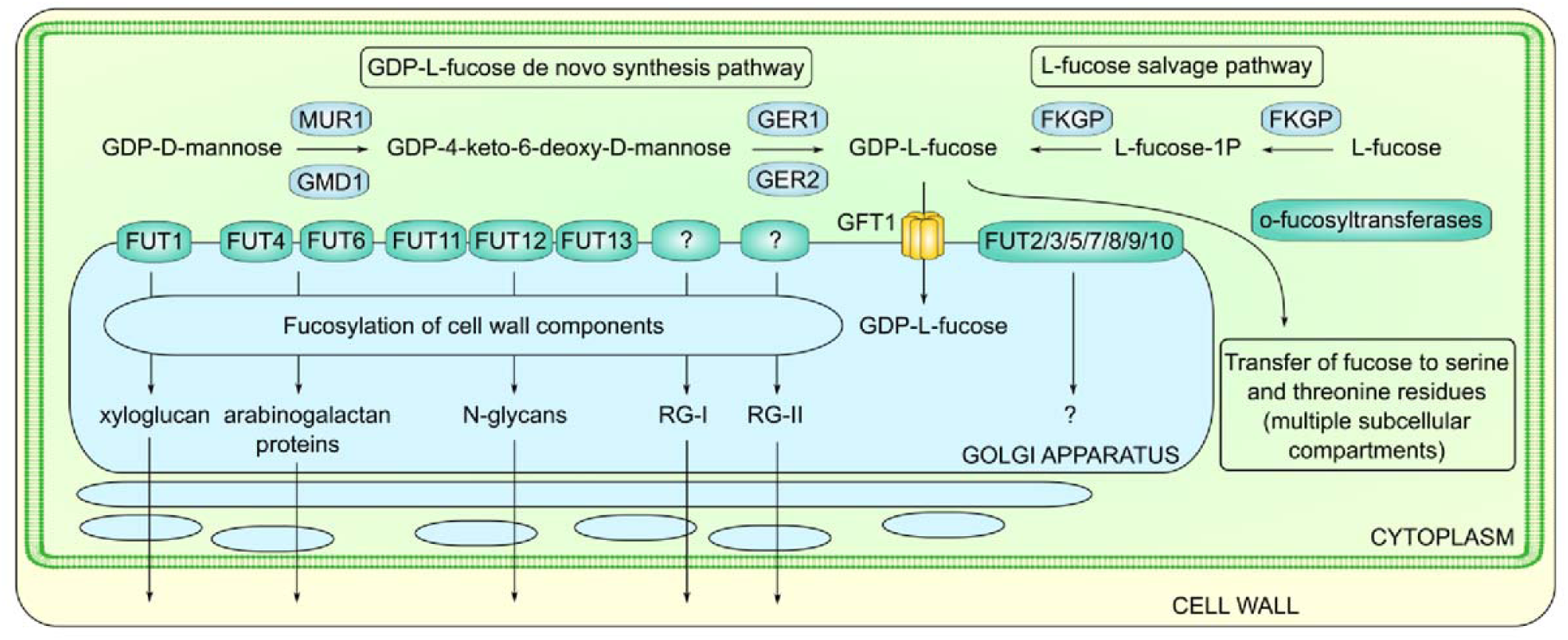
**Synthesis and metabolism of GDP-L-fucose in *Arabidopsis thaliana*.**

Plants lacking MUR1 exhibit dwarfism, reduced apical dominance, brittle stems (Reiter et al., 1993), short root phenotype (Van Hengel and Roberts, 2002), altered lignin structure and inflorescence stem development (Voxeur et al., 2017), and sensitivity to freezing (Panter et al., 2019) and pathogens (Zhang et al., 2019). The product of the MUR1-catalyzed reaction, GDP-4-keto-6-deoxy-D-mannose, serves as a substrate for the GDP-4-keto-6-deoxymannose-3,5-epimerase-4-reductases GER1 (Bonin and Reiter, 2000; Nakayama et al., 2003) and GER2 (Rhomberg et al., 2006) that complete the synthesis of GDP-L-Fuc (Figure 1). Another pathway that leads to synthesis of GDP-L-Fuc (the L-fucose salvage pathway) involves a single bifunctional enzyme L-FUCOKINASE/GDP-L-FUCOSE PYROPHOSPHORYLASE (FKGP) that converts L-Fuc to GDP-L-Fuc (Figure 1; Kotake et al., 2008). In contrast to *mur1*, the *fkgp* mutants exhibit normal growth phenotype and cell wall composition, suggesting that the L-fucose salvage pathway has a minor role in Arabidopsis (Kotake et al., 2008).

Following synthesis in the cytoplasm, GDP-L-Fuc is transported into the Golgi lumen by the GDP-FUCOSE TRANSPORTER1 (GFT1; Figure 1; Rautengarten et al., 2016). Knock-out mutants of GFT1 are not viable, and GFT1 knockdown plants exhibit semi-lethal phenotypes similar to those of *mur1* mutants (Rautengarten et al., 2016). Within the Golgi lumen, GDP-L-Fuc serves as a substrate for fucosyltransferases (FUT) that fucosylate components of the cell wall (Figure 1). There are 13 fucosyltransferases (FUT1 – FUT13) in Arabidopsis (Sarria et al., 2001; Wilson et al., 2001), and according to the current state of knowledge, all are either confirmed or expected to localize to the Golgi membrane with the catalytic domain facing the lumen (Sarria et al., 2001; Strasser, 2016). FUTs add L-Fuc residues onto molecules such as xyloglucan (FUT1/MUR2)(Perrin et al., 1999; Vanzin et al., 2002), arabinogalactan proteins (FUT4, FUT6; Wu et al., 2010; Liang et al., 2013; Tryfona et al., 2014) and N-linked glycans (FUT11/FUCTA, FUT12/FUCTB, FUT13/FUCTC; Leonard et al., 2002; Strasser et al., 2004). Furthermore, L-Fuc is found in RG-I and RG-II, although the FUTs involved in the synthesis of these pectins are not known. In terms of quantity, the majority of L-Fuc is found in RG-I (Anderson et al., 2012). Importantly, the dwarf phenotype of *mur1* mutants has been previously attributed to deficiency in boron-dependent dimerization of RG-II (O’Neill et al., 2001). In *mur1*, RG-II L-Fuc residues are replaced by L-galactose (Zablackis et al., 1996) which leads to an approximately 50% decrease in RG-II dimer formation (O’Neill et al., 2001) possibly caused by RG-II chain A truncation (Pabst et al., 2013).

To understand processes controlling stomatal movements, O_3_ can be used as an apoplastic ROS donor to stimulate stomatal closure (Kollist et al., 2007; Vahisalu et al., 2010). Ozone enters plants through stomata and subsequently decomposes to various ROS that further provoke active production of ROS by plant cells (Vainonen and Kangasjärvi, 2014). Plants deficient in O_3_-induced stomatal closure receive high doses of O_3_ that trigger formation of visible hypersensitive response-like lesions. These lesions are easy to score and allow the identification of stomatal mutants in forward genetic approaches (Overmyer et al., 2000). Previously, such a genetic screen led to the identification of several proteins involved in stomatal closure, i.e., SLOW ANION CHANNEL1 (SLAC1; Vahisalu et al., 2008), the receptor-like pseudokinase GUARD CELL HYDROGEN PEROXIDE-RESISTANT1 (GHR1) that participates in activation of SLAC1 (Sierla et al., 2018), and HIGH TEMPERATURE1 (HT1) kinase which acts in high CO_2_-induced guard cell signaling as an inhibitor of SLAC1 activation (Hõrak et al., 2016). Here we present an O_3_ exposure-based forward genetic screen specifically aimed at the identification of novel components involved in stomatal closure. We report the isolation of 76 novel mutants, and 130 new mutants of established stomatal regulators. In particular, we describe the identification of the MUR1 mutant and show that synthesis and import of GDP-L-fucose into the Golgi lumen play an important role in stomatal closure. Our results are consistent with the hypothesis that the stomatal deficiencies observed in *mur1* mutants are due to altered mechanical properties of the guard cell walls, and we propose that reduced dimerization of rhamnogalactouronan-II contributes to the impaired stomatal function observed in *mur1* mutants.

## RESULTS

### A forward genetic screen identifies novel regulators of guard cell signaling

To fill the gaps in known guard cell signaling networks, we have performed forward genetic screen based on an O_3_-sensitivity (Figure 2). This screen is fully independent from our previous screen (Overmyer et al., 2000). A total of 125,000 seeds of Arabidopsis line pGC1:YC3.6 (Yang et al., 2008), expressing yellow cameleon 3.6 (YC3.6; Nagai et al., 2004), a biosensor probe that allows for imaging of intracellular Ca^2+^, were treated with ethyl methanesulfonate (EMS). Later, 380,000 M2 plants were exposed to O_3_. Individual plants displaying visible O_3_-damage were then subjected to thermal imaging and water loss assay (see Materials and Methods). The progeny of the most prominent mutants (approximately 3,200 lines) was subjected to a secondary screen utilizing the same screening methods. Lines for which at least one of the phenotypes was clearly confirmed (551 lines) were then analyzed with a whole-rosette gas exchange system (Kollist et al., 2007) to investigate stomatal responses to a variety of stimuli inducing stomatal closure i.e. apoplastic ROS (delivered by means of O_3_-fumigation), 5 µM ABA spray, high CO_2_ and reduced air humidity which is often termed as vapor pressure deficit (VPD; Merilo et al., 2018). Lines demonstrating lack, or impairment of stomatal responses to at least a single stimulus (206 lines) were subjected to targeted sequencing of genomic regions encoding 22 well-established stomatal regulators, hereafter referred to as “*usual suspects*”, to avoid potential re-discoveries (Figure 2). The list of *usual suspects* included ion channels, ABA biosynthesis enzymes, protein kinases, phosphatases, and other proteins where the corresponding mutant lines are known to exhibit impaired stomatal closure and/or higher stomatal conductance (see Supplementary Table 1 for a full list). Approximately 60% of the tested lines (130) had mutations in coding regions of at least one *usual suspect*. Most frequently mutations were identified within *AHA1*, *GHR1* (Sierla et al., 2018), *MITOGEN-ACTIVATED PROTEIN KINASE12* (*MPK12*) and *MORE AXILLARY BRANCHES2* (*MAX2*) coding sequences (Figure 2). Among the 76 mutants with no mutations in *usual suspects*, the most frequently observed stomatal phenotypes were impaired responses to elevated CO_2_ (48 out of 76) and high loss of water from detached leaves (46 out of 76). Notably, the majority of newly identified mutants were affected in stomatal responses to more than one stimulus (Figure 2). With the exception of new alleles of HT1 and AHA1 that will be published elsewhere, all new alleles of the usual suspects are listed in (Supplementary Data Set 1). Due to the high number of mutant lines, we were not able to perform allelism tests for all new alleles of the *usual suspects*. However, published results (Sierla et al., 2018), as well as currently ongoing experiments, indicate that in most of the lines the observed phenotypes are linked to the mutations in the tested genes. Therefore, only the lines with no mutations in the *usual suspects* were retained for further analysis.

**Figure 2.**
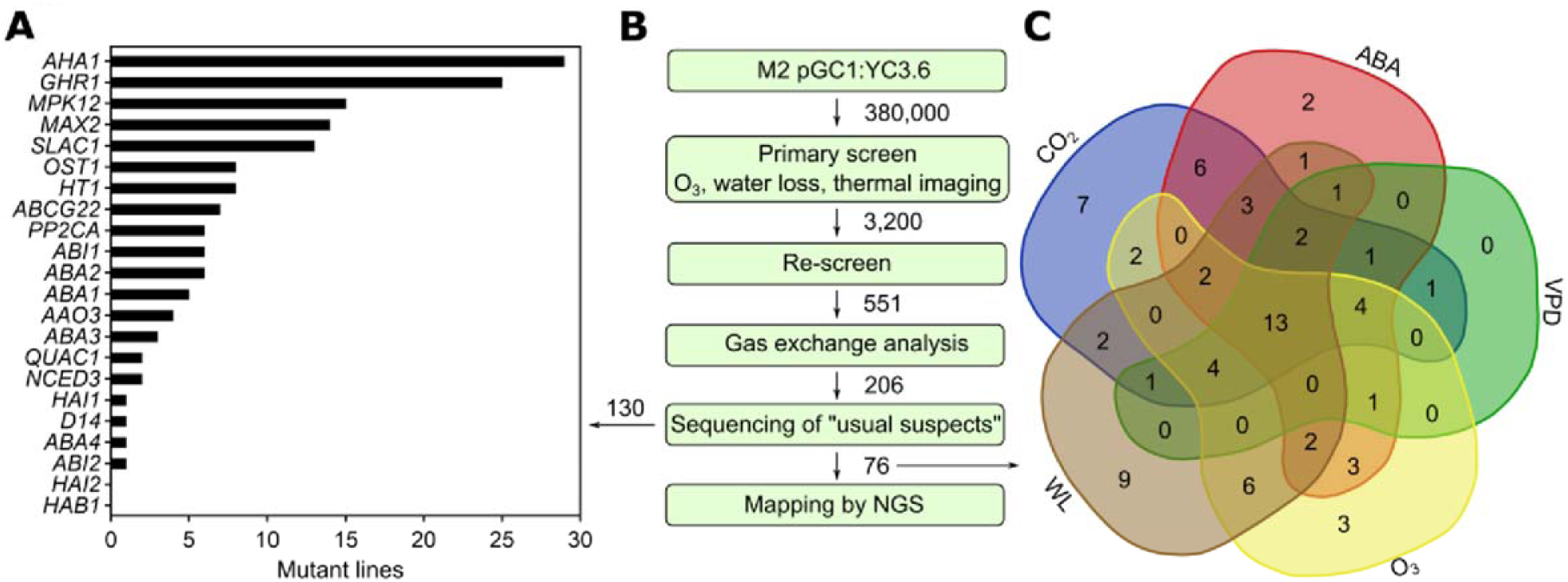
**Ozone sensitivity-based forward genetic screen to identify novel regulators of stomatal closure.** **(A)** The amount of mutant lines of known stomatal regulators identified during candidate gene sequencing. **(B)** Scheme of the screening procedure. **(C)** Whole-rosette gas exchange analysis of novel mutants impaired in stomatal closure. Numbers indicate the amount of mutants impaired in stomatal closure to indicated stimuli.

### Mapping of a novel mutant impaired in stomatal closure identifies MUR1

From the screen described above, mutant *T7-9* exhibited higher water loss from detached leaves (water loss) compared to wild type (Figure 3A), and partially impaired stomatal responses to O_3_, high CO_2_ concentration, ABA, and darkness, while retaining the ability to close stomata in response to high VPD (Supplemental Figure 1). Throughout this paper, the water loss assay, also known as “mass loss of detached leaves, MLD” (Duursma et al., 2019) is used as a simple indicator of stomatal function (if the cuticle permeability is intact) and has proven reliable in our previous work on stomatal signaling (Vahisalu et al., 2008; Hõrak et al., 2016; Sierla et al., 2018). To identify the causative mutation in *T7-9*, we applied the SHOREmap backcross pipeline (Hartwig et al., 2012). For this, *T7-9* was backcrossed to YC3.6. The resulting BC1_F1_ plants had a WT-like water loss indicating recessive inheritance. Approximately 21% of the BC1_F2_ plants (123 out of 593) exhibited increased water loss, indicating that the trait was determined by a single locus. Nuclear DNA from the 123 BC1_F2_ plants displaying the mutant phenotype was bulked and subjected to next generation sequencing (NGS). Analysis of the NGS data with marker frequency threshold set to 0.9 resulted in identification of five non-synonymous EMS-specific mutations enriched in BC1_F2_ plants displaying high water loss (Supplementary Table 2). All polymorphisms localized to the lower arm of chromosome 3 within a 17.0 - 19.1 Mb physical interval. Screening of the mutant lines for the five candidate genes revealed that three independent mutant lines: *mur1-1*, *mur1-2* (Reiter et al., 1993; Bonin et al., 1997) and *mur1-9* (SALK_057153, Supplementary Figure 2), carrying mutations within *AT3G51160* (Figure 3B), exhibited highly elevated water loss (Figure 3C). The mutant lines for the remaining candidate genes had WT-like water loss (Supplemental Figure 3). *AT3G51160* encodes GDP-mannose-4,6-dehydratase MURUS1 (GMD2, MUR1) which catalyzes the first step in *de novo* biosynthesis of GDP-L-Fucose (Figure 1; Bonin et al., 1997). Therefore, we investigated the level of GDP-L-Fuc in the *T7-9* mutant. Similarly to other *mur1* mutants, we were not able to detect this metabolite in *T7-9* suggesting the complete loss of MUR1 enzymatic activity (Figure 3D). Finally, an allelism test between *T7-9* and a *mur1-2* mutant revealed lack of complementation, confirming that the *T7-9* MUR1 E175K mutation (hereafter referred to as *mur1-10*) conferred its high water loss (Figure 3E).

**Figure 3.**
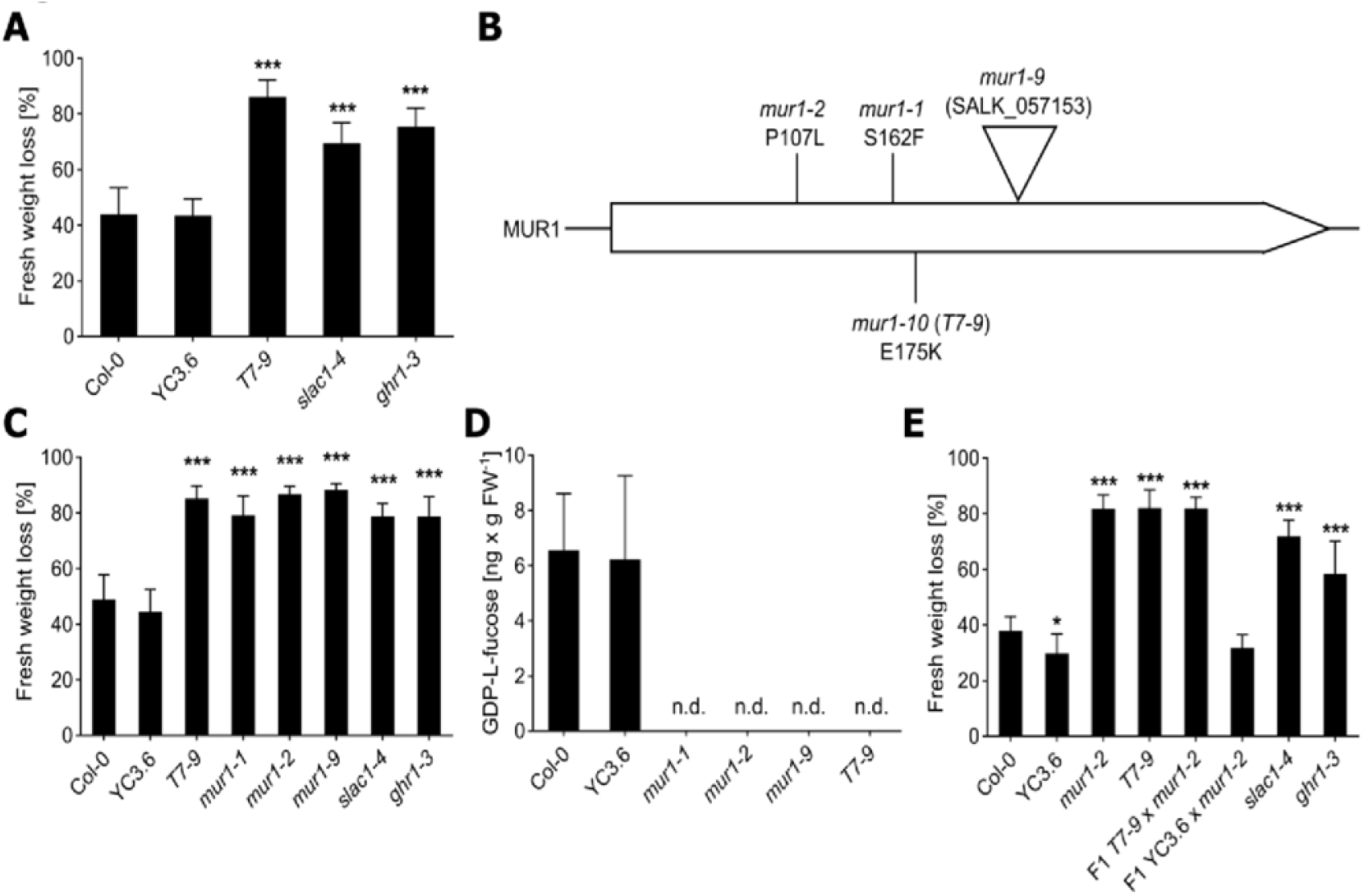
**Mapping of *T7-9* mutant.** **(A)** Leaf fresh weight loss of *T7-9* and control lines (YC3.6, Col-0, *slac1-4* and *ghr1-3*) recorded after 2h. Data bars represent means ± SD (n = 12 plants). **(B)** Positions of mutations in *mur1* mutants used in this study. **(C)** Leaf fresh weight loss of *T7-9*, independent *mur1* mutants (*mur1-1, mur1-2, mur1-9*) and control lines (YC3.6, Col-0, *slac1-4* and *ghr1-3*) recorded after 2h. Data bars represent means ± SD (n = 12 plants). **(A, C)** Asterisks denote statistical differences (*** p < 0.001) to respective control lines (Col-0 or YC3.6) according to one-way ANOVA followed by Sidak’s post-hoc test. **(D)** GDP-L-fucose content in *T7-9*, *mur1* mutants and respective control lines measured by UPLC-MS. Data bars represent means ± SD (n = 4-5 plants); n.d., not detected. **(E)** Leaf fresh weight loss of *T7-9*, *mur1-2*, F1 *T7-9 x mur1-2*, F1 YC3.6 x *mur1-2* and control lines recorded after 2h. Data bars represent means ± SD (n = 9-12 plants). Asterisks denote statistical differences (* p < 0.05; ** p < 0.01; *** p < 0.001) to Col-0 according to one-way ANOVA followed by Dunnett’s post-hoc test. **(A, C, E)** Experiments were repeated three times with similar results. Results of the representative experiments are shown.

### MUR1 is involved in stomatal development

As multiple phenotypes observed in *mur1* mutants have been attributed to abnormal cell wall composition (Reiter et al., 1993; O’Neill et al., 2001; Van Hengel and Roberts, 2002), we first focused on characterizing *mur1* stomata and cuticle development. No phenotypes related to cuticle permeability were detected in any of the four *mur1* mutants by means of dye-exclusion assay (Supplemental Figure 4). To assess stomatal development, we performed scanning electron microscopy-based examination of abaxial epidermis of *mur1-1* and *mur1-2* cotyledons. In agreement with an earlier report (Zeng et al., 2011), no consistent phenotype related to stomatal density was observed in *mur1* mutants. The stomatal density in *mur1-2* mutant was higher than in *mur1-1,* which had a similar stomatal density to Col-0 (Supplementary Figure 5A). The average size of stomatal pores was moderately increased in the *mur1-1* mutant, while in *mur1-2* there were no significant differences compared to Col-0 (Supplementary Figure 5B). Notably, in both *mur1* mutants we observed a larger variability in size of stomatal pores (Supplementary Figure 5C) and the biggest stomatal complexes (2-8% of stomata, Supplementary Figure 5D) were irregular in shape and often obstructed (Supplementary Figure 5E). Moreover, as observed before (Zeng et al., 2011; Zhang et al., 2019), a fraction of *mur1* stomata exhibited aberrant structure of the outer cuticular ledge. In the most severe cases (3-17% of stomata, Supplementary Figure 5D) the outer stomatal ledges completely sealed the stomatal pores (Supplementary Figure 5F) similarly as in mutants lacking FUSED OUTER CUTICULAR LEDGE1 (FOCL1), a proline-rich protein necessary for formation of outer cuticular ledge (Hunt et al., 2017). Overall, despite the differences in stomata size and morphology, the whole plant steady state stomatal conductance of the *mur1* mutants did not differ significantly from that of the wild type plants (Supplementary Figure 1, Supplementary Figure 6, Figure 4I).

**Figure 4.**
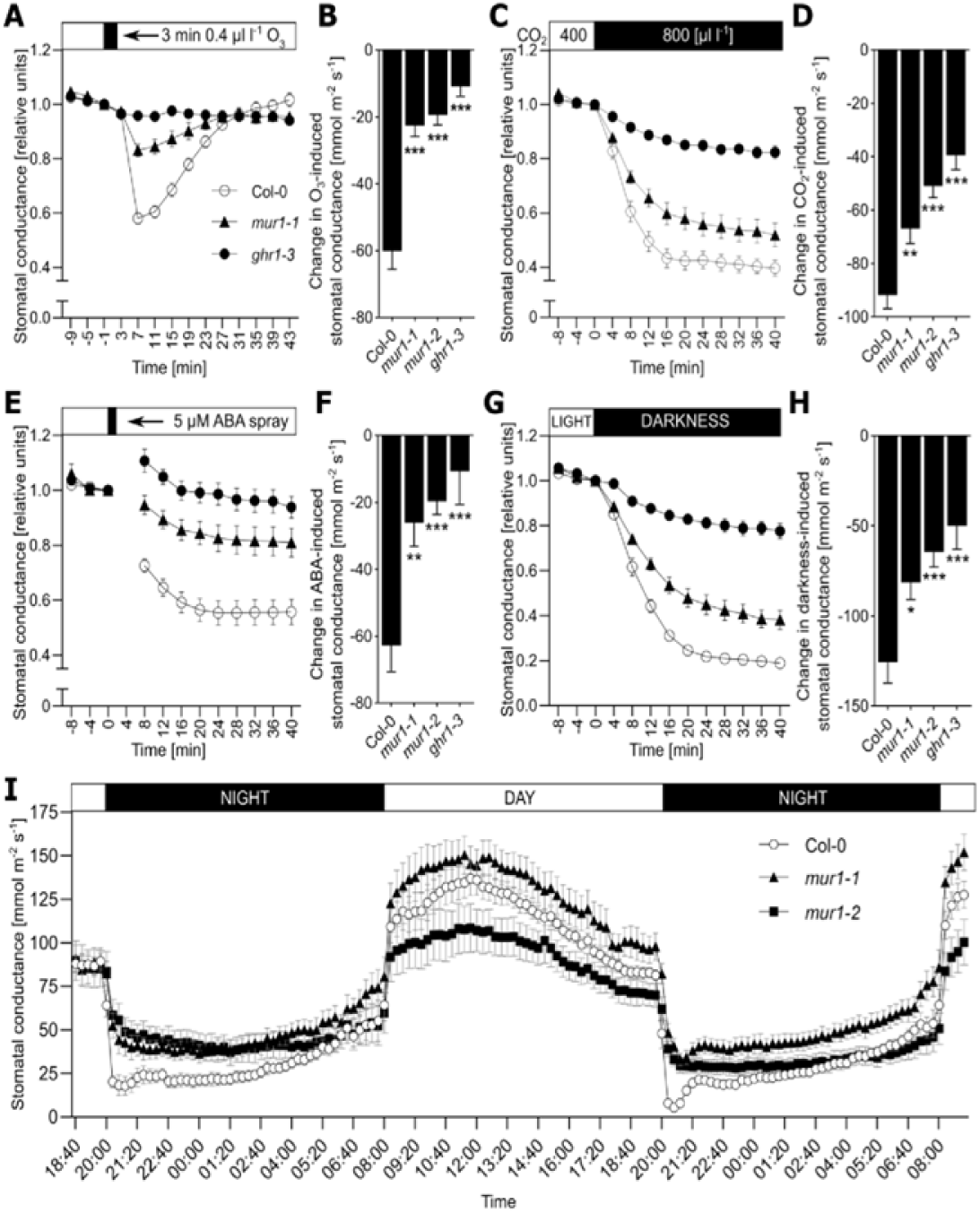
**Characterization of *mur1* stomatal phenotypes.** **(A-H)** Stomatal responses of *mur1* mutants to stomata-closing stimuli. The changes in stomatal conductance are shown in relative and absolute values calculated from the data presented in Supplementary Figure 6. **(A, C, E, G)** Time course of relative stomatal conductance (normalized to the last time point before the treatment) of 3- to 4-week-old *mur1-1*, Col-0, and *ghr1-3* plants in response to **(A)** O_3_ pulse, **(C)** elevated CO_2_, **(E)** ABA spray and **(G)** darkness. The indicated treatments were applied at t = 0 and whole-rosette stomatal conductance was recorded. Data points represent means ± SEM; n = 7-10 **(A)**, 10-12 **(C)**, 9-11 **(E)**, 6-11 **(G)** plants analyzed in two (A, E, G) or three (C) independent experiments. **(B, D, F, H)** Changes in stomatal conductance of Col-0, *mur1-1*, *mur1-2* and *ghr1-3* in response to **(B)** O_3_ pulse, **(D)** elevated CO_2_, **(F)** ABA spray and **(H)** darkness. Values were calculated by subtracting the initial stomatal conductance at t = 0 **(D, F, H)** or t = −1 **(B)** from the stomatal conductance at **(B)** t = 7 min, **(D, F, H)** t = 40 min. Data bars represent means ± SEM; n = 7-10 **(B)**, 10-12 **(D)**, 9-11 **(F)**, 6-11 **(H)** plants. Asterisks denote statistical differences to Col-0 (* p < 0.05; ** p < 0.01; *** p < 0.001) according to one-way ANOVA followed by Dunnett’s post-hoc test. **(I)** Diurnal changes in whole-rosette stomatal conductance of *mur1-1*, *mur1-2* and Col-0 plants. Data points represent means ± SEM; n = 8 plants analyzed in 3 experiments. Each experiment was started at the same time of the day and measurements were recorded for 45 h.

### MUR1 is required for stomatal closure

To further characterize the stomatal function in *mur1* mutants, we subjected them to a variety of treatments provoking stomatal movements and followed the time-resolved whole-rosette stomatal conductance (Kollist et al., 2007). Gas exchange measurements were performed for *mur1-1* and *mur1-2* mutants (Reiter et al., 1993; Bonin et al., 1997). For the treatments inducing stomatal closure, either *ghr1-3* (Sierla et al., 2018) or *ost1-3* (Yoshida et al., 2002) were used as non-responsive controls, while in stomata opening assays *ht1-2* (Hashimoto et al., 2006) was used. Across all gas-exchange assays, the stomatal responses of *mur1* mutants were consistent (Supplementary Figures 6 and 7), therefore, representative results obtained for *mur1-1* allele are shown (Figure 4). As observed earlier in *T7-9*, (Supplementary Figure 1) plants lacking MUR1 were impaired in the rapid transient decrease of stomatal conductance in response to a three-minute O_3_ pulse (Figure 4A-B, Supplementary Figure 6A), that otherwise induces a rapid decrease in Col-0 plants (Kollist et al., 2007; Vahisalu et al., 2010; Sierla et al., 2018). This suggests that the activity of MUR1 is required for rapid stomatal movements in response to ROS. Further, we observed impaired stomatal closure upon treatment with elevated CO_2_ concentration (800 µl l^-1^), 5 µM ABA spray or application of darkness during the light period (Figure 4C-H, Supplementary Figure 6B-D). Similarly, during diurnal light/dark cycles, the transition to darkness induced a rapid drop in stomatal conductance of Col-0 plants while the *mur1* mutants exhibited a much less pronounced response, and maintained higher stomatal conductance during the darkness period (Figure 4I). In contrast to these observations, as seen earlier in *T7-9* (Supplementary Figure 1), the stomatal responses of *mur1* mutants to high VPD were not significantly different from those of Col-0 plants (Supplementary Figure 6E-F). The stimuli provoking stomatal opening, such as exposure to low CO_2_ concentration (400 → 100 µl l^-1^) or increase in light intensity (150 → 500 µmol m^-2^ s^-1^), did not show any differences between *mur1* mutants and the wild type (Supplementary Figure 7) suggesting that MUR1 activity is not required for stomatal opening. Taken together, our data indicate that MUR1 activity is required for normal stomatal closure in response to O_3_, high CO_2_ concentration, ABA and darkness, but not for stomatal opening.

### Import of GDP-L-fucose into the Golgi apparatus is necessary for stomatal function

To further study the role of MUR1 in stomatal closure, we investigated stomatal function in mutants impaired in the subsequent steps of the GDP-L-Fuc synthesis and transport (Figure 1). For this, two T-DNA insertion mutants of GER1, (*ger1-2*, *ger1-3*) as well as GER2 (*ger2-1*, *ger2-2*) were isolated (Supplementary Figure 2B, C) and subjected to a water loss assay. None of the tested *ger* mutants exhibited elevated water loss, suggesting functional redundancy (Figure 5A). Similarly, the water loss of *fkgp* mutants (*fkgp-1, fkgp-2*; Kotake et al., 2008) was similar to that of Col-0 (Figure 5A) indicating that the L-fucose salvage pathway has little impact on stomatal closure. To further explore the possible redundancy between GER1 and GER2, we attempted to generate a *ger1 ger2* double mutant. To this end, *ger1-2* and *ger1-3* were crossed with *ger2-1* and *ger2-2*. However, no double homozygous plants were found in any of the four F2 families. Similarly, the F3 progeny of F2 plants homozygous for *ger1* but heterozygous for *ger2* alleles (*ger1-2^-/-^ ger2-1^+/-^*; *ger1-2^-/-^ ger2-2^+/-^*; *ger1-3^-/-^ ger2-1^+/-^*; *ger1-3^-/-^ ger2-2^+/-^*) did not contain double mutants. In every case, the observed segregation of *ger2* alleles (*ger2^+/+^* : *ger2^+/-^* : *ger2^-/-^*) within the F3 families was 1 : 1 : 0 (Supplementary Table 3). We therefore inspected both pollen viability and embryo development in F2 *ger1^-/-^ ger2^+/-^* plants but did not find any abnormalities. Hence, the inability to generate *ger1 ger2* mutants might be related to defects in fertilization with *ger1^-^ ger2^-^* pollen. Thus, we focused on characterization of the GDP-L-Fuc transporter GFT1.

**Figure 5.**
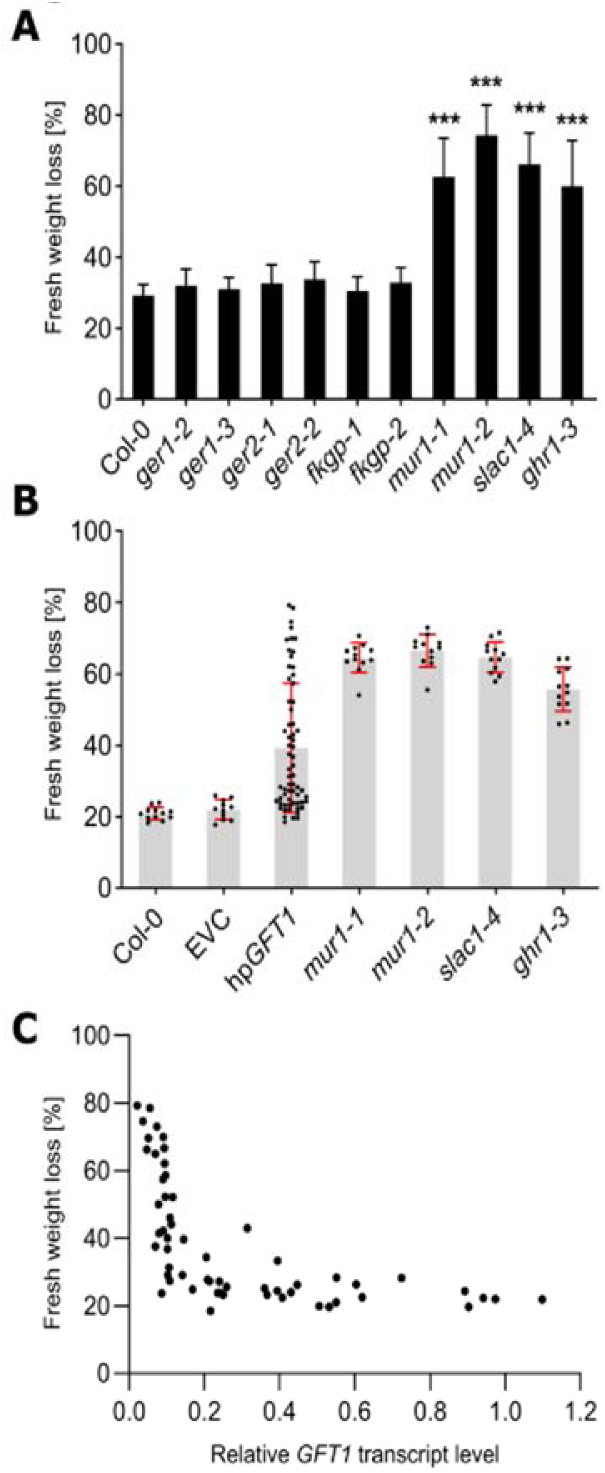
**Stomatal function in mutants affected in GDP-L-fucose metabolism.** **(A)** Leaf fresh weight loss of mutants of genes encoding enzymes involved in synthesis of GDP-L-fucose, and control lines (Col-0, *mur1-1*, *mur1-2*, *slac1-4* and *ghr1-3*), recorded after 2h. Data bars represent means ± SD (n = 13-16 plants). Asterisks denote statistical differences (*** p < 0.001) to Col-0 according to one-way ANOVA followed by Dunnett’s post-hoc test. Experiment was repeated three times with similar results. Results of the representative experiment are shown. **(B)** Leaf fresh weight loss of 64 independent hp*GFT1* T1 plants, and control lines (Col-0, EVC – empty-vector control, *mur1-1*, *mur1-2*, *slac1-4* and *ghr1-3*), recorded after 2h. Data points represented values obtained for separate plants. Data bars represent means ± SD (for control lines n = 11-12 plants). **(C)** Correlation between residual *GFT1* transcript level and fresh weight loss observed in 58 independent hp*GFT1* T1 plants. Each dot represents values obtained for an independent hp*GFT1* T1 plant.

Loss-of-function mutants of GFT1 are not viable, for that reason we utilized hairpin RNAi knockdown plants (hp*GFT1*; (Rautengarten et al., 2016). A total of 66 independent hp*GFT1* T1 plants were selected and transplanted to soil. As described before, the hp*GFT1* plants exhibited varying growth phenotypes, i.e., reduced projected rosette area, short petioles, and wavy leaves (Supplementary Figure 8A) that correlated with the residual *GFT1* transcript level and cell wall L-Fuc content (Rautengarten et al., 2016). For each of the T1 hp*GFT1* plant that survived in soil (64 plants) we measured the projected rosette area and loss of water from detached leaves, as well as the *GFT1* transcript level (58 plants). The hp*GFT1* T1 plants exhibited varying water loss (Figure 5B) and residual *GFT1* transcript level (Supplementary Figure 8B), and displayed a clear negative correlation between these two traits (Figure 5C). Similarly, the projected rosette area was also negatively correlated with the water loss (Supplementary Figure 8C). The majority of hp*GFT1* plants with *GFT1* transcript level lower than 10% of that observed in the empty vector control (EVC) displayed water loss comparable to that of *mur1* mutants (Figure 5C). Together, our data indicate that the import of GDP-L-Fuc into the Golgi lumen is important for stomatal closure.

### Lack of MUR1 affects mechanical properties of guard cell walls

To investigate the genetic interactions between MUR1 and proteins regulating canonical stomata closure pathways we crossed *mur1-1* and *mur1-2* mutants to *slac1-4* (Vahisalu et al., 2008), *aba2-11* (González-Guzmán et al., 2002), *ost2-2D* (Merlot et al., 2007), *ost1-3* (Yoshida et al., 2002) and *ghr1-3* (Sierla et al., 2018) and assayed the stomatal function of the double mutants via water loss assay. In every double mutant an additive effect of combining two mutations (Figure 6A) indicated that the phenotypes observed in *mur1* plants were independent from the canonical guard cell signaling pathways. Because of the additive effects (Figure 6A), the previously documented role of MUR1 in cell wall development (Reiter et al., 1993; Reiter et al., 1997; O’Neill et al., 2001), as well as the general deficiency in responses to stomata-closing stimuli observed in *mur1* mutants (Figure 4, Supplementary Figure 1, Supplementary Figure 6), we investigated the mechanical properties of *mur1* guard cell walls with atomic force microscopy (Carter et al., 2017). The patterning of the apparent modulus (E_a_) in the stomatal complexes of *mur1* mutants was comparable to that of the control lines. However, comparison of the absolute E_a_ values derived from the AFM scans indicated that *mur1* mutants had significantly stiffer subsidiary cells and this difference was even more pronounced when values obtained for guard cell walls were compared (Figure 6B). Thus, taken together, our data indicate that the stomatal phenotypes observed in *mur1* mutants are most likely related to the altered mechanical properties of the stomatal complexes.

**Figure 6.**
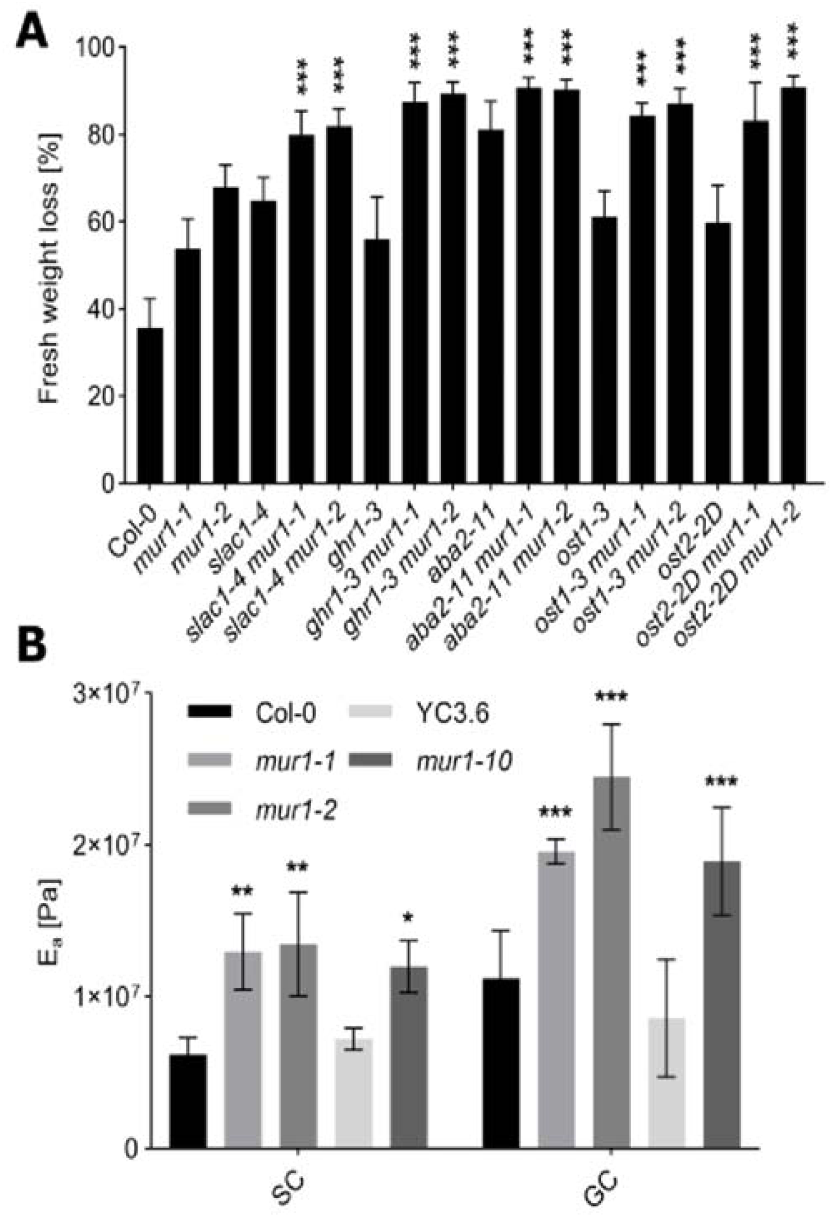
**Stomatal phenotypes observed in *mur1* mutants are independent from canonical guard cell signaling.** **(A)** Leaf fresh weight loss of double mutants obtained after crossing *slac1-4*, *ghr1-3*, *aba2-11*, *ost1-3* and *ost2-2D* with *mur1-1* and *mur1-2* recorded after 1h. Data bars represent means ± SD (n = 13-16 plants). Asterisks denote statistical differences (*** p < 0.001) to respective single mutant lines (*slac1-4*, *ghr1-3*, *aba2-11*, *ost1-3* and *ost2-2D*) according to one-way ANOVA followed by Sidak’s post-hoc test. Experiment was repeated three times with similar results. Results of the representative experiment are shown. **(B)** Average apparent Young’s modulus (E_a_) values derived from AFM scans of subsidiary cells (SC) and guard cells (GC) of control (Col-0, YC3.6) and *mur1* plants. Bars represent means ± SD (n = 2-3 plants, 2 stomata per plant). Data analyzed with two-way repeated measures ANOVA with “genotype” and “cell type” as factors, followed by Tukey’s post-hoc test. Asterisks denote statistical differences (* p < 0.05, ** p < 0.01, *** p < 0.001) to respective control lines (Col-0, YC3.6) observed within a cell type.

### *mur1* stomatal phenotypes are not linked to xyloglucan structure

Xyloglucan has been previously linked to stomatal movements (Rui and Anderson, 2016). To investigate whether the phenotypes observed in *mur1* mutants might be related to the lack of xyloglucan fucosylation, we analyzed the stomatal function in mutants lacking FUT1/MUR2, the major xyloglucan fucosyltransferase (Figure 1; Vanzin et al., 2002), and KATAMARI1/MURUS3 (KAM1, MUR3) a xyloglucan galactosyltransferase (Madson et al., 2003). The cell wall fucose content in *mur2-1, mur3-1* and *mur3-2* mutants is reduced by approximately 50 % (Reiter et al., 1997). Unlike MUR3 EMS mutants (*mur3-1*, *mur3-2*) the T-DNA insertion mutants *mur3-3* and *mur3-7* exhibit a dwarf rosette phenotype (Tamura et al., 2005; Kong et al., 2015) which can be rescued by removing the activity of XYLOSYLTRANSFERASE1 (XXT1) and XXT2 leading to plants devoid of xyloglucan (Kong et al., 2015). Therefore, we assessed the stomatal function of *mur2-1*, all four *mur3* mutants, as well as a *mur3-3 xxt1 xxt2* triple mutant (Kong et al., 2015) and an *xxt1 xxt2* double mutant (Cavalier et al., 2008). The water loss of *mur2-1* was not significantly different from that observed for Col-0 control, indicating that the lack of xyloglucan fucosylation does not affect the stomatal closure process (Figure 7A). All *mur3* mutants exhibited a moderately elevated water loss, however, much lower than that of *mur1* mutants. The water loss of xyloglucan-deficient *xxt1 xxt2* double mutant was similar to that of Col-0 (Figure 7A). Therefore, we concluded that the stomatal phenotypes observed in *mur1* mutants were not linked to defects in XyG fucosylation.

**Figure 7.**
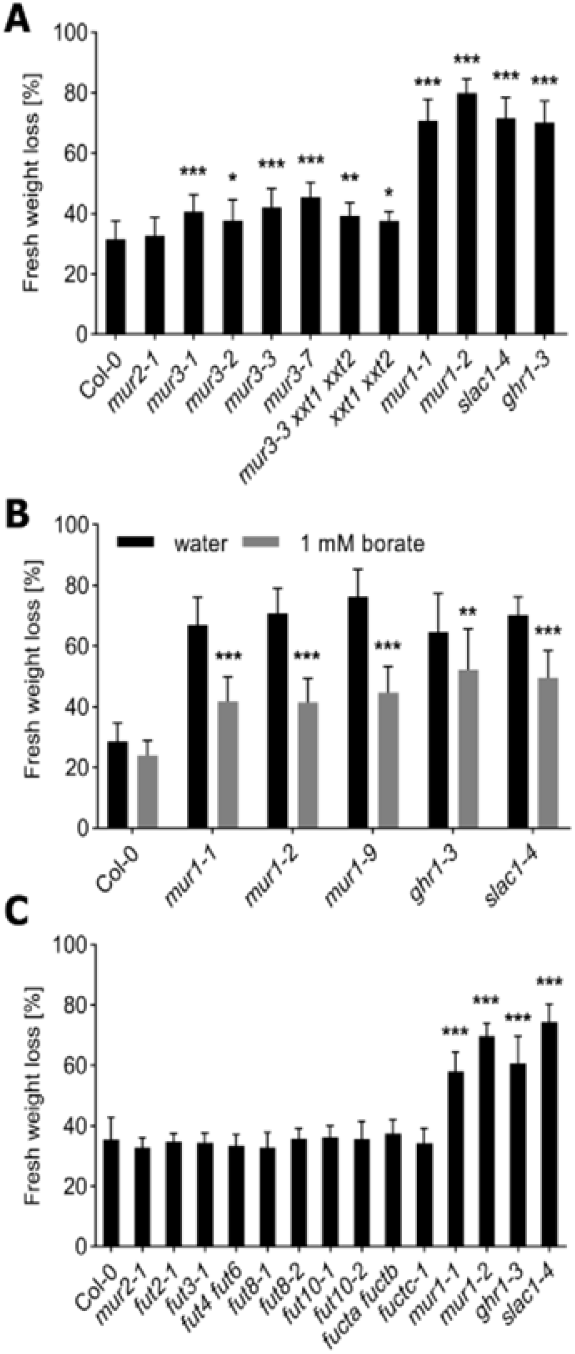
**Screen for fucose-containing cell wall components affecting stomatal function.** **(A)** Leaf fresh-weight loss of mutants affected in fucosylation and synthesis of xyloglucan, recorded after 2h. Values obtained for Col-0, *mur1-1*, *mur1-2*, *slac1-4* and *ghr1-3* are provided for reference. Data bars represent means ± SD (n = 15-16 plants). **(B)** The effect of borate supplementation on the leaf fresh-weight loss of *mur1* mutants and control lines (Col-0, *ghr1-3* and *slac1-4*). Bars represent means ± SD (n = 14-16 plants). Data analyzed with two-way ANOVA with “genotype” and “borate concentration” as factors, followed by Sidak’s post-hoc test. Asterisks denote statistical significances (** p < 0.01, *** p < 0.001) of treatment effect within each genotype. Experiment was repeated five times with similar results. Results of the representative experiment are shown. **(C)** Leaf fresh-weight loss of mutants deficient in fucosyltransferases. Values obtained for Col-0, *mur1-1*, *mur1-2*, *slac1-4* and *ghr1-3* are provided for reference. **(A, C)** Experiments were repeated at least 3 times with similar results. Results of the representative experiments are shown. Asterisks denote statistical differences (* p < 0.05, ** p < 0.01, *** p < 0.001) to Col-0 according to one-way ANOVA followed by Dunnett’s post-hoc test.

### *mur1* stomatal phenotypes are related to pectin structure

To investigate whether the stomatal phenotypes of *mur1* were related to the dimerization of RG-II, we grew *mur1* mutants in soil supplemented with 1 mM borate. Such treatments were previously shown to compensate for the deficiency in RG-II crosslinking (O’Neill et al., 2001). We observed a significant decrease of water loss of *mur1* mutants grown in the presence of 1 mM borate as compared to the control conditions (Figure 7B). A similar trend was observed in Col-0, *slac1-4* and *ghr1-3* mutants, albeit to a much lower extent. To validate this finding, we utilized plants lacking boron transporter REQUIRES HIGH BORON1 (BOR1) required for boron xylem loading (Noguchi et al., 1997; Takano et al., 2002) as previously the impaired uptake of boron was demonstrated to affect the dimerization of RG-II (Miwa et al., 2013; Panter et al., 2019). Lack of BOR1 leads to impaired expansion of rosette leaves which can be rescued by increase of B concentration in the growth medium (Noguchi et al., 1997). We found that a soil–grown *bor1-3* mutant (Kasai et al., 2011) exhibited high water loss which could be reverted by supplementing the soil with 50 µM borate, while lower concentrations (10 µM and 20 µM) had no effect (Supplementary Figure 9). Thus, the data supported the hypothesis that the phenotypes observed in *mur1* plants were related to a deficiency in RG-II crosslinking.

To further investigate the role of RG-II in stomatal closure, we analyzed the stomatal function of mutants lacking other enzymes involved in RG-II biosynthesis. The structure of RG-II is highly complex and in Arabidopsis only few enzymes involved in its synthesis have been identified (Funakawa and Miwa, 2015). Thus far, apart from enzymes involved in synthesis or transport of GDP-L-Fuc, the following proteins have been shown to play a role in RG-II synthesis: RHAMNOGALACTURONAN XYLOSYLTRANSFERASE1 (RGXT1), RGXT2 (Egelund et al., 2006), RGXT3 (Egelund et al., 2008), RGXT4/MGP4 (Liu et al., 2011) catalyzing the transfer of D-xylose onto L-fucose, KDO-8-P SYNTHASE1 (*At*KDSA1) and *At*KDSA2, required for synthesis of 3-deoxy-d-manno-octulosonate (KDO; Matsuura et al., 2003; Delmas et al., 2008); and GOLGI GDP-L-GALACTOSE TRANSPORTER1 (GGLT1) required for import of GDP-L-galactose into the Golgi apparatus (Sechet et al., 2018). Deficiency in RGXT4 or KDSA activity leads to defective pollen tube formation, preventing the generation of mutant lines (Delmas et al., 2008; Liu et al., 2011). Therefore, we investigated the water loss of plants deficient in RGXT1 (*rgxt1-1*), RGXT2 (*rgxt2-1*; Egelund et al., 2006), KDSA1 (*AtkdsA1-S*), KDSA2 (*AtkdsA2-S*; Delmas et al., 2008) and three GGLT1 RNAi knockdown lines (Sechet et al., 2018); but no elevated water loss was observed in any of the tested lines (Supplementary Figure 10). It should be noted that, under our growth conditions, none of the tested mutants phenocopied the *mur1* rosette phenotype, which suggests functional redundancy between the members of the respective gene families (Delmas et al., 2008) or different structural consequences for the cell wall caused by the alterations of RG-II composition in the respective mutants.

In addition, we investigated whether *mur1* stomatal phenotypes might be related to lack of fucosylation of other cell wall components. For this, we measured water loss of mutants deficient in FUT4 and FUT6 (*fut4 fut6*; Tryfona et al., 2014), FUT11 and FUT12 (*fucta fuctb*; Strasser et al., 2004), FUT13 (*fuctc-1*; Rips et al., 2017) as well as other fucosyltransferases with yet unidentified targets for which the mutant lines were available: *fut2-1*, *fut3-1*, *fut8-1*, *fut8-2*, *fut10-1* and *fut10-2* (Supplementary Figure 11). We did not detect *FUT9* transcript in 2-week-old seedlings, and at the time of the analysis no mutant lines for FUT5 and FUT7 were available, therefore FUT5, FUT7 and FUT9 were not included in these experiments. The water loss of *fut4 fut6*, *fucta fuctb* and *fuctc-1* mutants was comparable to that of Col-0 (Figure 7C), suggesting that *mur1* stomatal deficiencies are not related to lack of fucosylation of arabinogalactan proteins or N-linked glycans. Similarly, mutants of the remaining fucosyltransferases did not show elevated water loss, which likely reflects a high degree of functional redundancy within the FUT family.

## DISCUSSION

### Forward genetic screen identifies new components of guard cell signaling network

Stomatal movements are coordinated by multiple signaling pathways that converge to regulate the activity of guard cell tonoplast and plasma membrane ion channels. Over the last three decades, many components of guard cell signaling networks have been identified (Hedrich, 2012; Sierla et al., 2016; Ehonen et al., 2019; Lawson and Matthews, 2020), however multiple key components await characterization. Here we describe a forward genetic screen that aims to saturate the gene network controlling stomatal closure. Our results indicate that the current gene network is far from complete as we identified 76 mutants affected in stomatal closure (Figure 2) that do not represent mutations in any of the 22 well-established stomatal regulators. The majority of the novel mutants were impaired in stomatal closure to several stimuli, however mutants deficient in responses to a single stimulus e.g. apoplastic ROS generated by O_3_ exposure, or CO_2_ were also identified (Figure 2). We expect that future characterization of the newly identified mutants will provide insight into global, and stimuli-specific regulation of stomatal closure. As expected, apart from identifying novel mutants, our screening strategy yielded 130 new mutants of genes encoding known stomatal regulators (Supplementary Data Set 1). Mutations in *AHA1*, *GHR1*, *MPK12* and *MAX2* (Figure 2) were the most prevalent, which might emphasize the significance of these genes in stomatal closure or the importance of intact protein sequence for the whole-protein function. New alleles of the *usual suspects* described here can act as a useful resource for detailed studies of their molecular function. Recently we used 10 novel alleles of *GHR1* to dissect the role of specific mutations on its function and stability (Sierla et al., 2018). Currently, we are investigating the newly identified mutants of *AHA1* and *HT1* to get a deeper insight into their mode of action. An unexpected finding was the identification of multiple novel alleles of *MAX2* (Supplementary Data Set 1). MAX2 is mostly recognized for its role in strigolactone signaling (Waters et al., 2017) and has been previously implicated in guard cell functions (Ha et al., 2014; Piisilä et al., 2015) that are likely independent from SLs signaling (Kalliola et al., 2020). However, our data suggests that the role of MAX2 in stomatal closure might be more central than previously realized. In conclusion, our work provides genetic resources that enable further studies of established stomatal regulators. Furthermore, mapping of newly identified mutants will contribute to saturation of the genetic landscape of stomatal regulation.

### Synthesis and import of GDP-L-fucose into the Golgi is necessary for stomatal closure

At the organellar/cellular level, the flux of water through the guard cell tonoplast and plasma membranes results in changes in volume and surface area of guard cells (Franks et al., 2001; Shope et al., 2003; Meckel et al., 2007) and their central vacuoles (Diekmann et al., 1993; Gao et al., 2005). Stomatal opening involves an increase in guard cell turgor (Franks et al., 2001), volume, and surface area that in turn lead to an increase in guard cell length (Meckel et al., 2007). Contrary to stomatal opening, stomatal closure involves guard cell shrinking. However, it is not entirely understood how the repetitive, and fast changes in guard cell wall surface area are executed during stomatal movements. According to current understanding, this process relies on the flexibility of the cell wall matrix which is determined during guard cell morphogenesis (Rui et al., 2018), and is presumably promoted by the turgor pressure of the subsidiary cells. Data available thus far point towards the key role of the pectin network in controlling the cell wall flexibility that enables stomatal movements, and results presented in this study support this hypothesis. The first mutant characterized from the screen described here corresponds to MUR1 - an enzyme catalyzing the first step in *de novo* GDP-L-Fuc synthesis pathway (Bonin et al., 1997). Plants lacking GDP-L-Fuc exhibited high loss of water from detached leaves and impaired responses to apoplastic ROS, ABA, darkness and high CO_2_ concentration (Figure 4, Supplementary Figures 1 and 6). Elevated water loss was also observed in plants impaired in import of GDP-L-Fuc into Golgi apparatus (Figure 5B-C). Therefore, we conclude that not just the synthesis, but also the import of GDP-L-Fuc into the Golgi lumen is necessary for stomatal closure. Recently, *mur1* mutants were also shown to lack stomatal responses to treatment with *Pseudomonas syringae* (*Pst* DC3118) and salicylic acid (Zhang et al., 2019). The stomatal defects observed in *mur1* appear independent from canonical guard cell signaling pathways (Figure 6A) and instead stem from defects in cellular metabolism.

The Golgi apparatus (GA) serves as a hub for synthesis of pectic polysaccharides which are later exported to the apoplastic space (Caffall and Mohnen, 2009). Accordingly, the fucosylation of cell wall polysaccharides, AGPs and N-linked glycans is thought to occur in the GA (Figure 1; Chou et al., 2015; Strasser, 2016) and the majority of fucose is incorporated into the cell wall via the GA-derived vesicles (Anderson et al., 2012). We found that mutants of fucosyltransferases responsible for synthesis of AGPs, N-linked glycans and xyloglucan did not exhibit elevated water loss (Figure 7 A,C), therefore we excluded the possibility that phenotypes observed in *mur1* might be linked to impaired fucosylation of these cell wall components. The partial reversion of the *mur1* water loss phenotype by borate (Figure 7B), and high water loss observed in *bor1-3* mutant (Supplementary Figure 9), suggests that the phenotype is linked either directly, or indirectly, to the structure and dimerization of RG-II. Earlier it was found that *mur1* mutants, even when grown under high borate conditions, had 74-78 % of RG-II in dimer form while nearly all RG-II (95%) was dimerized in Col-0 (O’Neill et al., 2001). Therefore, partial restoration of *mur1* stomatal function might be related to incomplete RG-II dimerization even after borate supplementation.

A complementary explanation for the observed phenotype might be related to shortening of the RG-II chain A observed in *mur1* plants (Pabst et al., 2013). As RG-II and HG share the same backbone and are covalently cross-linked (Caffall and Mohnen, 2009; Harholt et al., 2010) the shortening of the RG-II chain A might have similar consequences as digestion of RG-I arabinan side chains (as observed previously by Jones et al., (2003)). According to this scenario, the lack of GDP-L-Fuc would promote crosslinking of HG and render the guard cell walls stiffer and, thus, less responsive to changes in turgor pressure. This hypothesis is supported by our AFM data as we found that *mur1* mutants have significantly stiffer epidermal cell walls than wild type plants (Figure 6B). It is noteworthy, that mutants with highly decreased HG content, *quasimodo1* (*qua1*; Bouton et al., 2002) and *qua2* (Mouille et al., 2007) exhibit morphological phenotypes related to loss of cell adhesion which are not a direct consequence of decreased HG content (Verger et al., 2016). In addition to impaired development, both mutants exhibit high loss of water from detached leaves (Bouton et al., 2002; Krupková et al., 2007) however, it is yet unknown whether this phenotype is related to impaired stomatal closure.

One argument against the hypothesis of increased HG crosslinking comes from the observation of the stomatal opening process in *mur1* mutants. In contrast to stimuli inducing stomatal closure, *mur1* mutants responded properly to stomatal opening cues such as increase in light intensity and decrease in CO_2_ concentration (Supplementary Figure 7), while Jones et al., (2003) observed impaired stomatal opening after the arabinanase treatment. However, a similar phenotype to that reported here, i.e. impaired closure and normal opening, was observed in plants lacking POLYGALACTURONASE INVOLVED IN EXPANSION3 (PGX3; Rui et al., 2017). Polygalacturonases constitute a large group of pectin-hydrolyzing enzymes (Yang et al., 2018). PGX3 is expressed in expanding tissues and guard cells where it controls the abundance and molecular mass of HG. PGX3 deficiency led to increased crosslinking of HG and impaired stomatal responses to ABA and darkness while the light- or fusicoccin-induced stomatal opening was not affected (Rui et al., 2017). On the basis of this phenotype Rui et al., (2017) proposed a model in which the loosening of pectin structure is necessary for stomatal closure, which is supported by the results of our study.

The normal stomatal opening observed in *mur1* and *pgx3* mutants might be explained by the relatively low severity of phenotypes observed in these two mutants. Jones et al., (2003) reported that treatment with arabinanase completely inhibited stomatal closure, while slower and “stepwise” stomatal closure was observed in *mur1* and *pgx3* mutants, respectively. The digestion of arabinan via exogenous enzyme treatment in epidermal strips likely affects the structure of the guard cell pectin network to a much greater extent than that observed in the intact leaves of *mur1* and *pgx3* mutants, leading to a stomatal opening phenotype.

Because of the observed changes in mechanical properties of guard cell walls, and the independence of *mur1* stomatal phenotypes from canonical guard cell signaling pathways, we hypothesize that the impaired function of *mur1* stomata is linked to a relatively stiffer epidermis. Since stomatal movements occur in the context of a mechanical environment determined by the subsidiary cells it is possible that the increased stiffness of the subsidiary cells also contributes to the *mur1* phenotypes.

### Alternative hypotheses and future challenges

Since *mur1* mutants exhibit a radical reduction in availability of GDP-L-Fuc, additional factors may contribute to impaired function of *mur1* stomata. For example, a fraction of stomatal pores of *mur1* plants were sealed with outer cuticular ledges (Supplementary Figure 5). A similar but more severe phenotype, which was also associated with impaired stomatal closure in response to ABA, has been observed in the *focl1* mutant (Hunt et al., 2017). The precise molecular function of FOCL1 is not known, however, it has been proposed to control the mechanical properties of guard cell walls and the formation of the cuticle-cell wall bond (Hunt et al., 2017). Thus, it is tempting to speculate that, while being independent from guard cell signaling, impaired *mur1* stomatal responses might be linked to those observed in *focl1*.

As the majority of cell wall L-fucose is localized in RG-I (Anderson et al., 2012), the lack of fucosylation of this cell wall component might also contribute to phenotypes observed in *mur1*. To our knowledge, the identity of fucosyltransferases responsible for incorporation of L-Fuc into RG-I and RG-II remains unknown. Despite screening of multiple fucosyltransferase mutants we were not able to reproduce the *mur1* phenotype (Figure 7C), which probably reflects functional redundancy within the FUT family. In support of this hypothesis, the coding sequences of multiple FUTs are highly similar and physically localized in groups (Sarria et al., 2001) which suggests that closely related FUTs arose during gene duplication and, thus, might function redundantly. Therefore, due to the current lack of genetic tools, we were unable to precisely pinpoint the cell wall components responsible for *mur1* stomatal phenotypes. Analysis of high-order mutant lines, or the application of a CRISPR- Cas/amiRNA strategy, might be suitable for identification of fucosyltransferases that fucosylate RG-I and RG-II.

Depletion of GDP-L-Fuc likely results also in defects in protein o-fucosylation. According to phylogenetic analysis of plant and metazoan o-fucosyltransferases, Arabidopsis POFUTs constitute a multigene family with nearly 40 members (Smith et al., 2018). Among them SPINDLY was shown to fucosylate and activate DELLA proteins (Zentella et al., 2017) and according to (Zhang et al., 2019), part of the pathogen susceptibility observed in *mur1* mutants can be explained by lack of SPINDLY-mediated protein o-fucosylation. However, we were not able to detect stomata-related phenotypes in *spy* mutant lines. Another two putative o-fucosyltransferases FRIABLE1 (FRB1) and ESMERALDA (ESMD) were shown to influence cell adhesion (Neumetzler et al., 2012; Verger et al., 2016). Similarly to *qua1* and *qua2*, plants lacking FRB1 exhibit loss of cell adhesion (Neumetzler et al., 2012). Strikingly, the introduction of the *esmd* mutation into *qua1*, *qua2* and *frb1* backgrounds restored their cell adhesion defects with no apparent changes in cell wall composition, implying the existence of a signaling pathway controlling cell adhesion (Verger et al., 2016). More research efforts are needed to investigate whether the above-discussed mutants are impaired in stomatal function.

In summary, our study reveals that the synthesis of GDP-L-fucose is necessary for stomatal closure and highlights the key role of fucose metabolism for stomatal closure. The interpretation of our data in the context of earlier observations/hypotheses (Jones et al., 2003; Amsbury et al., 2016; Rui et al., 2017) leads us to conclude that the impaired stomatal closure observed in *mur1* is linked to increased stiffness of guard cell walls which likely stems from enhanced crosslinking of HG caused by impaired structure of RG-II. While the information obtained from mutant lines reported here provides valuable indications as to which cell wall components are necessary for the execution of stomatal movements, precisely how wall structure is modified/reorganized upon perception of stomatal opening/closing stimuli to accommodate volume/pressure changes awaits future investigation.

## MATERIALS & METHODS

### Plant material and growth conditions

All *Arabidopsis thaliana* mutants used in this study were in the Col-0 genetic background. The following lines were obtained from Nottingham Arabidopsis Stock Center: *mur1-1, mur1-2* (Reiter et al., 1993; Bonin et al., 1997); *mur1-9* (SALK_057153); *ger1-2* (WiscDsLox_425G11), *ger1-3* (GK_296H02), *ger2-1* (GK_113G05), *ger2-2* (SALK_091781); *mur2-1* (Reiter et al., 1997; Vanzin et al., 2002); *ghr1-3* (GK_760C07; Sierla et al., 2018); *slac1-4* (SALK_137265; Vahisalu et al., 2008); *ost1-3* (SALK_008068; Yoshida et al., 2002); *mur3-7* (SALK_127057; Tamura et al., 2005); *AtkdsA1-S* (SALK_024867), *AtkdsA2-S* (SALK_066700; Delmas et al., 2008); *rgxt1-1* (SALK_073748), *rgxt2-1* (SALK_023883; Egelund et al., 2006), *bor1-3* (SALK_037312; Kasai et al., 2011),*fut2-1* (GK_320C07), *fut3-1* (SALK_045666), *fut8-1* (SALK_010965), *fut8-2* (WiscDsLox_449F06), *fut10-1* (WiscDsLox_432A01), *fut10-2* (SALK_020408), *fuctc-1* (SALK_067444; Rips et al., 2017). pGC1:YC3.6 line (Yang et al., 2008) was donated by Julian Schroeder. FKGP mutants *fkgp-1* (SALK_012400), *fkgp-2* (SALK_053913; Kotake et al., 2008) were donated by Toshihisa Kotake; *mur3-1*, *mur3-2* (Reiter et al., 1997), *mur3-3* (SALK_141953; Tamura et al., 2005), *xxt1* (SAIL_785_E02) *xxt2* (SALK_101308) double mutant (Cavalier et al., 2008), and *xxt1 xxt2 mur3-3* triple mutant (Kong et al., 2015) were donated by Malcolm A. O’Neill; *ost2-2D* mutant (Merlot et al., 2007) was donated by Jeffrey Leung; *aba2-11* mutant (González-Guzmán et al., 2002) was donated by Pedro L. Rodríguez; *fucTA* (SALK_087481) *fucTB* (SALK_063355) double mutant (Strasser et al., 2004) seeds were obtained from Richard Strasser; *fut4* (SAIL_284_B05) *fut6* (SALK_078357) double mutant seeds (Tryfona et al., 2014) were obtained from Paul Dupree; *snrk2.236* triple mutant seeds (Fujii and Zhu, 2009; Cui et al., 2016) were donated by Hiroaki Fujii and Kirk Overmyer; *ht1-2* mutant (Hashimoto et al., 2006) was donated by Koh Iba. All mutant lines were genotyped with gene specific primers (Supplementary Data Set 2). GGLT1 RNAi lines (Sechet et al., 2018), together with the corresponding empty-vector control, were donated by Julien Sechet and Jenny C. Mortimer; T1 seeds of hp*GFT1* and empty-vector control as well as *E.coli* and *Agrobacterium tumefaciens* strains carrying the hp*GFT1* construct in pART27 (Rautengarten et al., 2016) were obtained from Joshua L. Heazlewood. To generate hp*GFT1* lines, Col-0 plants were transformed via the floral-dip method with *Agrobacterium tumefaciens* strain AGL1 carrying the hp*GFT1* construct in pART27 vector (Rautengarten et al., 2016). T1 transformants were selected on half-strength Murashige & Skoog medium (Duchefa Biochemie) supplemented with 50 µg ml^-1^ kanamycin, in a controlled growth chambers (MLR-350, Sanyo) under 12 h light (130-160 µmol m^-2^ s^-1^)/12 h dark cycle, 22°C/18°C (day/night). After two weeks plants were transplanted to soil and grown for additional three weeks as described below. The same selection procedure has been applied to select for empty-vector control lines. Col-0, *mur1-1*, *mur1-2*, *slac1-4* and *ghr1-3* lines were grown in parallel, except the selection antibiotic was not included in the growth medium. The double mutants of *mur1-1* and *mur1-2* with *ost1-3*, *aba2-11*, *ghr1-3*, *slac1-4* and *ost2-2D* were generated by crossing *mur1-1* and *mur1-2* (pollen acceptors) with the respective pollen donor. Double mutants were identified in F2 generation by genotyping with gene-specific primers (Supplementary Data Set 2). An attempt to generate *ger1 ger2* double mutants was performed by crossing *ger1-2* and *ger1-3* (pollen acceptors) with *ger2-1* and *ger2-2* (pollen donors). Later, F2 populations were genotyped with gene-specific primers (Supplementary Data Set 2). F2 plants homozygous for *ger1* but heterozygous in *ger2* locus were allowed to self-pollinate and the F3 plants were genotyped as before to determine the segregation of *ger2* alleles.

Unless specified otherwise, seeds were suspended in 0.1% agarose solution, vernalized in the dark for two days at 4°C and sown on a 1:1 mixture of peat and vermiculite. Plants were grown in controlled growth rooms under 12 h light (200 µmol m^-2^ s^-1^)/12 h dark cycle, 22°C/18°C (day/night), 60%/70% relative humidity.

### Mutagenesis and screening procedure

Mutagenesis of Arabidopsis pGC1:YC3.6 (Yang et al., 2008) seeds was performed as described before (Kim et al., 2006). In the primary screen, M2 plants were grown for two weeks and exposed to 275-350 nl l^-1^ O_3_ for 6 h. Presence of O_3_-induced lesions was assessed visually, and sensitive plants were rescued. One to two weeks later, rosettes were imaged with an Optris PI450 thermal imager (Optris GmbH) and subjected to water loss assay as described below. M3 seeds were collected from the M2 plants that exhibited the most pronounced phenotypes i.e. severe O_3_ sensitivity, high loss of water and/or low leaf temperature. Later, to confirm the inheritance of the observed phenotypes, the M3 plants were exposed to 350 nl l^-1^ O_3_ for 6 h, and subjected to water loss assay. Approximately half of both, the primary and the secondary screen, was performed at the ExpoSCREEN facility (HMGU, Munich) and the remaining half at the plant growth facilities of the University of Helsinki. Lines for which at least one phenotype was reproduced, were selected for whole-rosette gas exchange-based phenotyping of stomatal function. Gas exchange measurements were performed on 2 - 4 plants per mutant line and in total 551 lines were analyzed for stomatal responses to O_3_ pulse (547 lines), elevated CO_2_ (550 lines), darkness (106 lines), VPD (498 lines) and ABA spray (482 lines). Stomatal conductance of 505 lines was tested.

Lines with defective stomatal responses to at least one stimulus were subjected to sequencing of “usual suspects” either by NGS-based sequencing of PCR amplicons obtained with the use of gene-specific primers (Supplementary Data Set 2) and Phusion DNA polymerase (ThermoFisher Scientific; ROUND 1 to 6), or whole-genome sequencing (ROUND 7 and 8). In the PCR-based approach, for each line, amplicons were pooled and subjected to NGS with the use of GS FLX+ system (Roche) or MiSeq system (Illumina) at DNA Sequencing and Genomics Laboratory, Institute of Biotechnology, University of Helsinki. Sequencing with the use of GS FLX+ system was performed exactly as described before (Sierla et al., 2018). For MiSeq-based sequencing, pooled amplicons were sheared (Bioruptor NGS, Diagenode) into approximately 500 bp fragments, end-repaired, A-tailed and ligated to truncated TruSeq adapters (Illumina). After purification a PCR reaction was performed to introduce full-length P5 adapter and indexed P7 adapter sequences. After PCR, all products were pooled, purified with AMPure XP (Agencourt, Beckman Coulter), size selected to 600-800 bp (as described above), and analyzed on a Fragment Analyzer (Advanced Analytical Technologies Inc., Ankeny, IA, USA). Paired-end sequencing were performed on a MiSeq using a 600 cycle kit v3 (Illumina). The obtained sequences were trimmed using Cutadapt (Martin, 2011) and assembled with SPAdes (Bankevich et al., 2012). In the whole genome sequencing approach (ROUND 7 and 8, performed at Institute for Molecular Medicine Finland, FIMM) leaf samples of approximately 10 plants per line were used for nuclear DNA isolation. Nextera Flex library preparation was performed using 50-100 ng of dsDNA according to Nextera DNA Flex Library Prep Reference Guide (Illumina, San Diego, CA, USA) with the following modifications: all reactions were performed in half of the normal volume and library normalization was done according to the concentration measured on LabChip GX Touch HT (PerkinElmer, USA). Later, 500-850 bp fragments were size selected from the pool using BluePippin (Sage Science, USA). In ROUND 7, sequencing was performed with Illumina NovaSeq 6000 system using S4 flow cell with lane divider (Illumina, San Diego, CA, USA). Read length for the paired-end run was 2×151 bp. In ROUND 8, sequencing was performed with the Illumina NovaSeq 6000 system using S1 flow cell with lane divider (Illumina, San Diego, CA, USA). Read length for the paired-end run was 2 x 101 bp. Following the data acquisition, reads were subjected to demultiplexing with Illumina bcl2fastq2.20 software.

Quality control of the reads was carried out using FastQC software (https://www.bioinformatics.babraham.ac.uk/projects/fastqc/), and the adapter removal and read trimming were performed with Trimmomatic v0.36 (Bolger et al., 2014). After removal of adaptor sequences, all bases were removed from the beginning and the end of the reads if their Phred quality score was below 20. Additionally, reads with mean Phred score <15 in a sliding window of 3 bases were clipped, and the reads with read length < 35 bp after trimming were removed. The trimmed reads were aligned to the TAIR10 version of the Arabidopsis genome using Bowtie 2 v2.2.9 (Langmead and Salzberg, 2012). Next, the sequence alignment data was converted to binary alignment file and sorted using Picard v2.5.0 (http://broadinstitute.github.io/picard/). The same toolbox was used for removing the read PCR duplicates and for adding read group information. Next, Genome Analysis Toolkit (GATK, https://gatk.broadinstitute.org) by Broad Institute was used to obtain genomic VCF files using HaplotypeCaller (Poplin et al., 2017), for combining the gVCF files, and finally for the joint calling of variants using GenotypeGVCFs. The called SNPs were annotated using snpEff v4.3t (Cingolani et al., 2012) and Arabidopsis gene annotation downloaded from TAIR on May 1^st^ 2018. The SNP calls in the list of usual suspects were then collected for manual inspection.

### Water loss measurement

Two middle age leaves of 3.5-week-old soil-grown plants were cut and dried abaxial side up at room temperature for 2 h, unless specified otherwise. Weight of leaves was determined before and after drying, and water loss was calculated as percentage of initial fresh weight loss.

### Quantification of GDP-L-fucose

Approximately 160 mg of frozen plant material was ground to powder and extracted first with 1 ml of 80% (v/v) methanol and then with 0.5 ml of 100% methanol. During each extraction, the homogenate was vortexed at 4 °C for 2 h, then centrifuged at 21,500 g for 5 min at 4 °C. The supernatants were combined and evaporated to dryness in vacuum (MiVac Duo concentrator, GeneVac Ltd, Ipswich, UK) and reconstituted in 50 µl of 50% (v/v) methanol. The GDP-L-fucose was quantified with UPLC-6500+ QTRAP/MS system (Sciex, CA) in negative (ESI-) multiple reaction monitoring (MRM) mode. Two MRM transitions for GDP-L-fucose (C_16_H_25_O_15_N_5_P_2_, molecular weight 589.34 g mol^-1^) were used: 588.0 → 442.0 for quantitative, and 588.0 → 424.0 for qualitative purposes. The chromatographic separation was performed in Waters BEH Amide column (100 mm × 2.1 mm, ø1.7 µm) at a flow rate of 0.4 ml min^-1^. Column compartment temperature of UPLC system (Sciex, CA) was set to 35 °C while the samples were kept at 10 °C. Initial chromatographic conditions were 15% buffer A (10 mM ammonium formate, pH 9.0) and 85% buffer B (acetonitrile). The initial conditions were held for 1 min and followed by a linear gradient to 60 % (A) in 6 min, then returned to initial conditions in 1 min and left to stabilize for 2 min. Analysis has been performed with the following parameters: declustering potential of −85, entrance potential of −10, collision cell exit potential of −36 and 40, respectively. The ion spray voltage was set at −4,500V, source temperature (TEM) at 425 °C, collision gas (CAD) was set to medium, curtain gas to 20, and source gas 1 (GS1) and 2 (GS2) were both set to 10. All data were acquired with Analyst 1.5.1 software (Sciex, CA) and GDP-L-fucose was quantified with MultiQuant 2.1 software (Sciex, CA) by utilizing standard calibration curve of GDP-L-fucose standard (Sigma Aldrich).

### Gas exchange analysis

The gas-exchange experiments were performed as described before (Sierla et al., 2018). Briefly, plants were grown in 4:3 peat : vermiculite mix under 12 h day (23°C) / 12 h night (18°C) regime at 70% relative humidity and 100-150 μmol m^-2^ s^-1^ light intensity. The stomatal function of 3 - 4 week-old plants was analyzed with the use of a custom-built gas-exchange device (Kollist et al., 2007). First, plants were inserted into the device and pre-incubated for about 1 h to stabilize the stomatal conductance. The experimental conditions in the chambers were as follows: ambient CO_2_ (∼400 µL L^- 1^), 150 µmol m^-2^ s^-1^ light, ∼70% relative air humidity, 24°C. Later, stimuli triggering stomatal movements were applied and changes in stomatal conductance were monitored in time. To induce stomatal closure O_3_ pulse (450 nl l^-1^ during 3 min), elevated CO_2_ (from 400 µl l^-1^ to 800 µl l^-1^), darkness, increase in VPD (reduction in relative air humidity (from 70% to ∼30-35%)) or 5 µM ABA spray were applied. Similarly, reduction of CO_2_ concentration (from 400 µl l^-1^ to 100 µl l^-1^) or increase of light intensity (from 150 μmol m^-2^ s^-1^ to 500 μmol m^-2^ s^-1^) were used to trigger stomatal opening. In order to study diurnal light/darkness responses, stomatal conductance was recorded for approx. 40 h under conditions that mimicked the growth day/night regime.

### Mapping by next-generation sequencing

Leaf tissue samples from the BC1_F2_ plants exhibiting the mutant phenotype were harvested in bulk and used for preparation of nuclear DNA. The nuclear DNA-enriched sample was sheared to 400 – 600 bp fragments (Bioruptor NGS), end-repaired, A-tailed and ligated to truncated TruSeq adapters (Illumina). Full-length P5 and indexed P7 adapter sequences were introduced by PCR. The library was purified and size selected to 500-700 bp fragments as described above. Paired-end sequencing (150 – 150 bp) of the obtained library was done on a NextSeq 500 sequencer (Illumina) to an approximately 50-fold genome coverage at DNA Sequencing and Genomics Laboratory, Institute of Biotechnology, University of Helsinki. Further, data were analyzed with SHORE v0.9.0 pipeline (Ossowski et al., 2008) implementing GenomeMapper v0.4.4s read alignment tool (Schneeberger et al., 2009) according to the procedure described earlier (Sun and Schneeberger, 2015). In parallel, the same procedure was performed for line YC3.6. Next, SHOREmap v3.2 (Sun and Schneeberger, 2015) was used to identify, prioritize and annotate mutations enriched in plants displaying mutant phenotype. This final step involved subtraction of all innate YC3.6 polymorphisms.

### Nuclear DNA enrichment

For nuclear DNA enrichment, plant material was ground in liquid nitrogen and 1 g of tissue powder was homogenized with 15 ml of HBM buffer (25 mM Tris-HCl pH 7.5, 440 mM sucrose, 10 mM MgCl_2_, 0.1% Triton X-100, 10mM *β*-mercaptoethanol, 2 mM spermine). The homogenate was filtered through Miracloth and centrifuged for 10 min at 1,075g, 4°C. Pellet was resuspended in 1 ml of NIB buffer [20 mM Tris-HCl pH 7.5, 250 mM sucrose, 5 mM MgCl_2_, 5 mM KCl, 0.1% (v/v) Triton X-100, 10 mM *β*-mercaptoethanol] and applied to a 15/50% (v/v) Percoll gradient in NIB buffer. Samples were centrifuged for 10 min at 3000g, 4°C and pelleted nuclei were resuspended in 0.5 ml NIB buffer and centrifuged again. Pellets were mixed with 750 µL of DNA extraction buffer [2 % (w/v) cetyltrimethylammonium bromide, 100 mM Tris-HCl pH 8.0, 20 mM EDTA pH 8.0, 1.4 M NaCl, 1 % (w/v) polyvinylpyrrolidone 40] and incubated for 30 min at 60°C. Next, samples were extracted with chloroform : isoamyl alcohol (24:1, v/v) and centrifuged for 10 min at 7,000g, 4°C. The water-soluble phase was collected, treated with RNase A (Merck), and DNA was precipitated with isopropanol at −20°C. Samples were centrifuged for 6 min at 13,000g, 4 °C, the pellet was washed twice in 70% (v/v) ethanol, air-dried, and resuspended in water. DNA concentration was measured with Qubit fluorometer (Thermo Fisher Scientific) according to manufacturer’s instructions.

### Expression Analysis by qPCR

For gene expression analysis of T-DNA insertion mutants, plants were grown vertically on half-strength Murashige & Skoog medium (Duchefa Biochemie) in controlled growth chambers (model MLR-350, Sanyo) under 12 h light (130-160 µmol m^-2^ s^-1^)/12 h dark cycle, 22°C/18°C (day/night). For every biological replicate (3 in total), approximately 10 whole two-week-old plants were pooled, frozen and ground in liquid nitrogen. Total RNA was extracted using a Genejet plant RNA isolation kit (Thermo Fisher Scientific). For analysis of *GFT1* transcript level, whole rosettes of T1 hp*GFT1* plants (minus two middle-aged leaves that were used for water loss assay) were frozen in liquid nitrogen and ground with a mortar and pestle. Total RNA was extracted as described above. Two micrograms of RNA were treated with DNaseI and reverse-transcribed using oligo-dT(20) priming with Maxima Reverse Transcriptase (RT) and Ribolock RNase inhibitor (Thermo Fisher Scientific) in a 31.5 µl volume. The reactions were diluted to the final volume of 100 µl, 1µl of which was used as template for PCR with 5x HOT FIREPol EvaGreen qPCR Mix Plus (no ROX; Solis Biodyne) with gene-specific primers (Supplemental Table 5). The PCR was performed on the CFX384 Real-Time System (Bio-Rad) with the following cycle conditions: 95°C 10 min, 60 cycles with 95°C 30 s, 60°C 10 s, 72°C 30 s, and ending with melting curve analysis. Col-0 cDNA dilution series was used to determine primer amplification efficiencies. Three reference genes were used for data normalization: *YELLOW-LEAF-SPECIFIC GENE8* (*YLS8, AT5G08290*), *TAP42 INTERACTING PROTEIN OF 41 KDA* (*TIP41, AT4G34270*), and *MONENSIN SENSITIVITY1 (MON1, AT2G28390*; Czechowski et al., 2005). Data were analyzed with qBase 3.0 (Biogazelle) and the transcript levels were related to those observed in respective control lines.

### Expression analysis by RT-PCR

For gene expression analysis of *fut* mutants plants were grown, and the cDNA was produced, as described above. PCR was performed with the use of FirePol polymerase (Solis Biodyne) and 1 µl of cDNA was used as a template. *PP2AA3* (*At1g13320*) transcript was used for reference. All primers are listed in Supplementary Data Set 2. The cycling conditions were as follows 95°C 3 min, 40 cycles with 95°C 30 s, 56°C 10 s, 72°C 90 s ending with 5 min final elongation at 72°C. PCR products were analyzed by agarose gel electrophoresis using ethidium bromide staining for imaging.

### Stomatal morphology and cuticle permeability

For scanning electron microscopy, cotyledons of 3-week-old soil-grown plants (1 cotyledon per plant, 4-6 plants per line per biological replicate, 3 biological replicates) were harvested and fixed for 16 h in a fixing buffer (2% paraformaldehyde, 2.5% glutaraldehyde, 0.1 M sodium cacodylate pH 7.4). Subsequently, samples were incubated for 1 h at room temperature in 0.1 M sodium cacodylate buffer pH 7.3 containing 1% (w/v) OsO_4_ and washed twice in distilled water. Later samples were dehydrated by 3 x 10 min washing in 50%, 70%, 96% and 100% (v/v) ethanol, subjected to critical point drying (Leica EM CPD300, Leica Microsystems GmbH), mounted on the aluminum stubs and coated with 5 nm layer of platinum (Quorum Q150TS, Quorum Technologies Ltd). Serial images (1,800x magnification, 10 % overlap) were taken with Quanta FEG 250 (Thermo Fisher Scientific) scanning electron microscope at the Electron Microscopy Unit, Institute of Biotechnology, University of Helsinki. Images were stitched in Fiji (Schindelin et al., 2012) implementing MIST plugin (Chalfoun et al., 2017). For each cotyledon from 1 to 2.5 mm^2^ area was analyzed, containing 100 - 350 stomatal complexes. Morphology of every stomata within the region of interest was assessed visually and assigned into one of the four categories: normal appearance, not determined, sealed (Supplementary Figure 5F), and obstructed (Supplementary Figure 5E). The size of stomatal pores was determined by measuring the length of the stomatal pore defined by the boarders of the outer cuticular ledge. Cuticle permeability assay was performed on middle-age leaves of 3-week-old soil-grown plants as described earlier (Cui et al., 2016) except 0.01% Tween 20 has been added to the staining solution. Mutants *aba2-11* (González-Guzmán et al., 2002) and *snrk2.236* (Fujii and Zhu, 2009) were used as controls with high cuticle permeability (Cui et al., 2016).

### Projected rosette area

Rosettes were photographed with Nikon D5100 camera equipped with AF-S Micro Nikkor 40 mm 1:2.8G objective (Nikon). The projected rosette area was determined by image analysis with ImageJ software (Schneider et al., 2012) equipped with Measure Rosette Area Tool (http://dev.mri.cnrs.fr/projects/imagej-macros/wiki/Measure_Rosette_Area_Tool).

### Atomic force microscopy

For AFM experiments, seeds were stratified for 7 days at 4°C, then grown in a 3:1 compost:perlite mix. Growth conditions were as follows: light intensity 170 μmol m^-2^ s^-1^, 12h day (21°C) / 12 h (17°C) night, 60% humidity. Leaves from approx. 21-day-old plants were analyzed as described before (Carter et al., 2017).

## SUPPLEMENTARY DATA

Supplementary Data Set 1. Mutations identified during candidate gene sequencing.

Supplementary Data Set 2. List of primers

## ACKNOWLEDGEMENTS

We thank Julian Schroeder, Toshihisa Kotake, Malcolm A. O’Neill, Jeffrey Leung, Pedro L. Rodríguez, Jenny C. Mortimer, Julien Sechet, Richard Strasser, Hiroaki Fujii, Kirk Overmyer, Koh Iba and Paul Dupree for providing seeds of multiple mutants used in this study. We thank Joshua Heazlewood and Berit Ebert for providing *Agrobacterium tumefaciens* strain carrying the hp*GFT1* construct, T1 seeds of hp*GFT1* and corresponding EVC. Anna Huusari, Konrad Łosiński, Jelena Odintsova and Samuli Lundström are acknowledged for excellent technical help. Leena Grönholm and Valtteri Lehtonen are acknowledged for maintaining the plant growth facilities. Personnel of the DNA sequencing and genomics laboratory is acknowledged for the technical assistance with NGS. Mervi Lindman is acknowledged for assistance with electron microscopy. We gratefully acknowledge Hans Lang for an excellent technical support during the ozone fumigation experiments at the EUS. We thank Andreas Albert and the technical staff of EUS for outstanding support during the experiment. Thanks to Mikael Brosché and Kirk Overmyer for helpful comments and discussions throughout the duration of the project. This work was supported by the University of Helsinki, Academy of Finland Centre of Excellence program (2014-2019), Academy of Finland post-doctoral fellowships (Decision 294580, C.W.; 266793, T.V.), University of Helsinki 3-year research grant (C.W., M.L.G., J.P.), EDUFI fellowship (M.L.G.), The Ella and Georg Ehrnrooth Foundation (M.S.), Finnish Cultural Foundation (M.S), Dora Plus Programme (Contract No. 36.9-6.1/1372, O.Z.), Estonian Research Council (PRG433, H.K.; PUT311, PRG719, D.Y.; Mobilitas Pluss postdoctoral researcher grant (Contract no. MOBJD291), A.K.P), European Regional Development Fund (Center of Excellence in Molecular Cell Engineering CEMCE, H.K.). The ozone exposure experiments performed at the HMGU were supported by Transnational Access program of the European Plant Phenotyping Network (EPPN, grant no. 284443) funded by the FP7 Research Infrastructures Programme of the European Union.

## AUTHOR CONTRIBUTIONS

J.K., conceived the project; C.W., T.V., D.Y., M.S., M.L.G., R.C., N.S., A.J.F., H.K., J.K., designed experiments, C.W., T.V., D.Y., M.S., O.Z., M.L.G., J.P., R.C., A.K.P., M.N., M.C., T.P., N.S., A.L., L.P., P.A., A.J.F., J.S., performed experiments and analyzed the data, J.D., D.E., J.B.W., provided technological solutions for large-scale O3 exposures; C.W., T.V., D.Y., M.S., A.J.F., H.K. and J.K. wrote the manuscript with comments from all co-authors.

## FIGURE LEGENDS

**Supplementary Figure 1.**
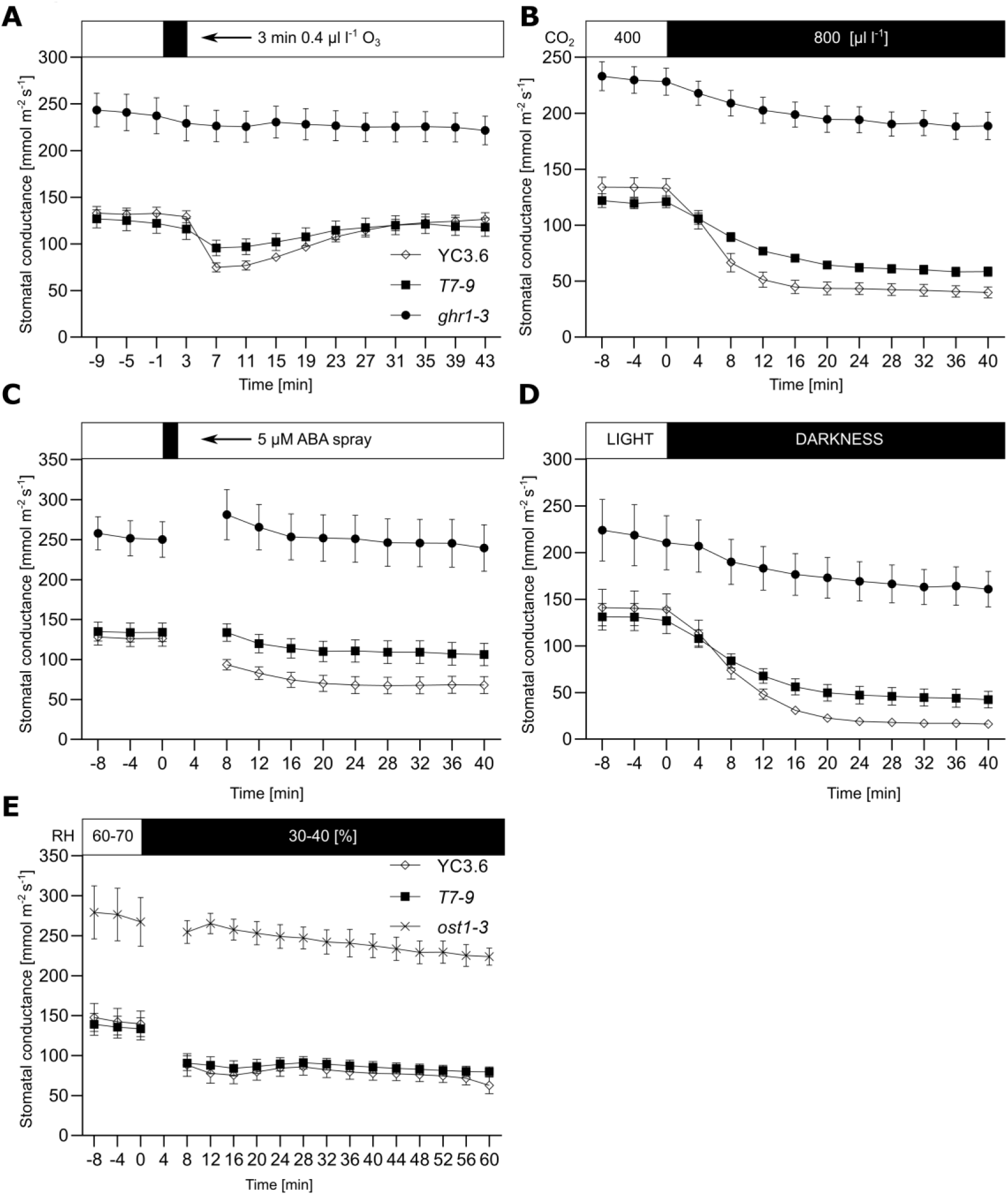
**Characterization of *T7-9* stomatal phenotypes.** Time course of whole-plant stomatal conductance of *T7-9*, YC3.6 and *ghr1-3* **(A-D)** or *ost1-3* **(E)** in response to **(A)** O_3_ pulse, **(B)** elevated CO_2_, **(C)** ABA spray, **(D)** darkness and **(E)** drop in relative air humidity (VPD increase from 1.01 ± 0.01 kPa to 2.04 ± 0.03 kPa (mean ± SE)). The indicated treatments were applied at t = 0 and whole-rosette stomatal conductance of 3- to 4-week-old plants was recorded. Data points represent means ± SEM; n = 8-9 **(A)**, 8-10 **(B)**, 9-10 **(C)**, 6-8 **(D)**, 3-12 **(E)** plants analyzed in 2 independent experiments.

**Supplementary Figure 2.**
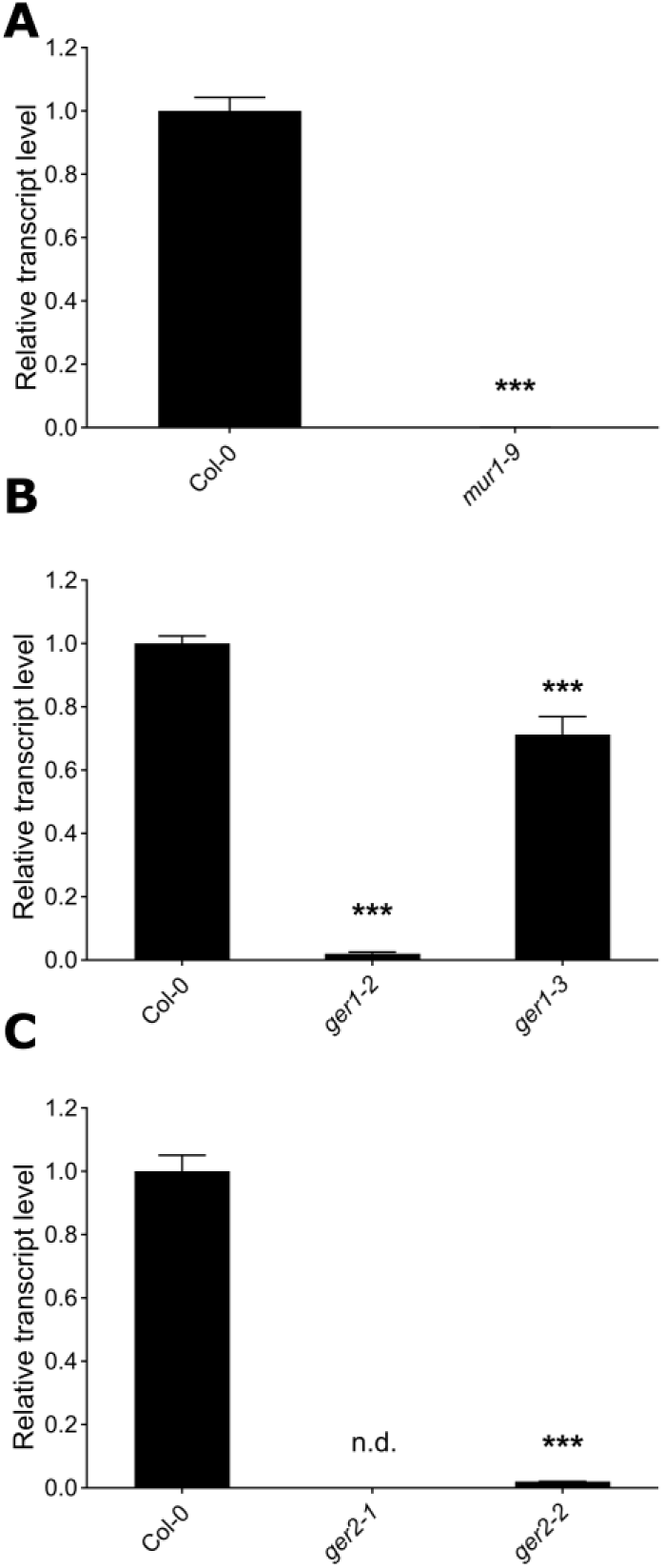
**Q-PCR analysis of *mur1-9*, *ger1* and *ger2* mutants.** **(A)** Relative *MUR1* transcript level in Col-0 and *mur1-9* plants. **(B)** Relative *GER1* transcript level in Col-0, *ger1-2* and *ger1-3* plants. **(C)** Relative *GER2* transcript level in Col-0, *ger2-1* and *ger2-2* plants. **(A-C)** Experiments performed on whole two-week-old plants grown in vitro on ½ x MS medium under 12 h light (120 – 160 µmol m^-2^ s-1)/12 h dark cycle at 22°C/18°C (day/night) temperature. Data bars represent means of three biological replicates ± SD. Asterisks indicate significant differences (*** p < 0.001) to Col-0 according to **(A)** Student’s *t*-test, **(B, C)** one-way ANOVA followed by Dunnett’s post-hoc test; n.d., not detected.

**Supplementary Figure 3.**
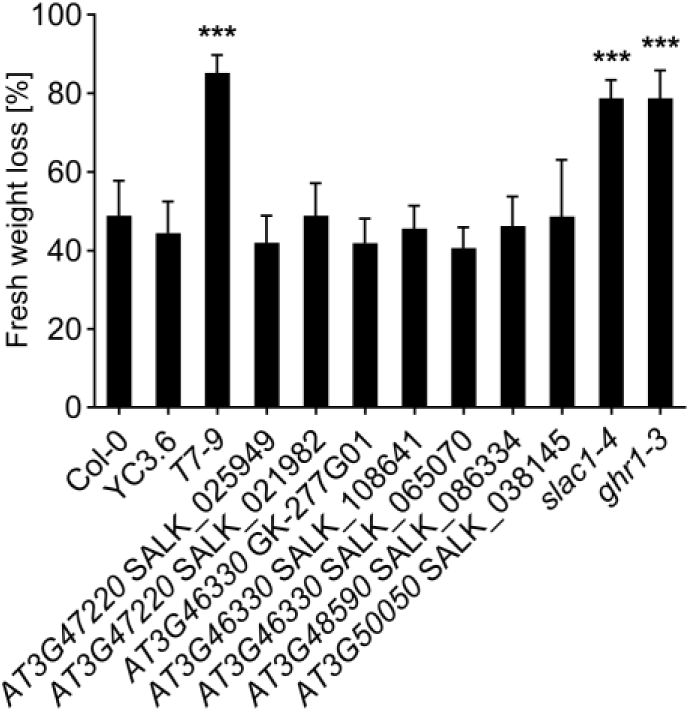
**Water loss-based screen of T-DNA insertion mutants of *T7-9* candidate genes.** Fresh weight loss of *T7-9*, control lines (YC3.6, Col-0, *slac1-4*, *ghr1-3*) and T-DNA insertion mutants of T7-9 candidate genes recorded after 2h. Data bars represent means ± SD (n = 12 plants). Asterisks denote statistical differences (*** p < 0.001) to respective control lines (Col-0 or YC3.6) according to one-way ANOVA followed by Sidak’s post-hoc test. This figure is an integral part of Figure 3C, values recorded for *T7-9* and control lines are provided for reference. Experiment was performed three times with similar results.

**Supplementary Figure 4.**
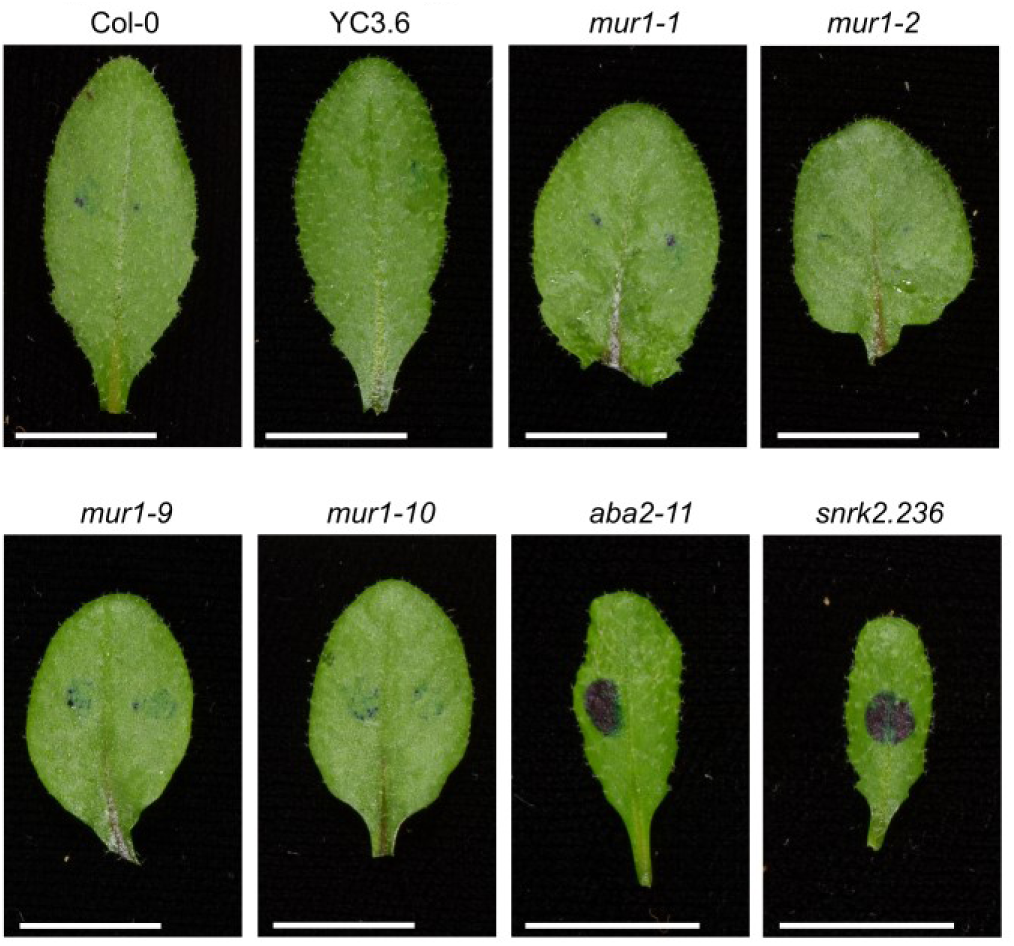
**Cuticle permeability of *mur1* mutants.** Representative photos of middle-aged leaves of 3.5 weeks-old *mur1* mutants, respective control lines and cuticule-deficient mutants (*aba2-11*, *snrk2.236*) subjected to dye exclusion assay. Leaves were stained with 5 µL drops of toluidine blue solution (0.05 % toluidine blue, 0.01% Tween 20) for 2h and rinsed with water. Dark blue staining indicates high cuticule permeability. Experiment was performed three times with similar results. Scale bar, 10 mm.

**Supplementary Figure 5.**
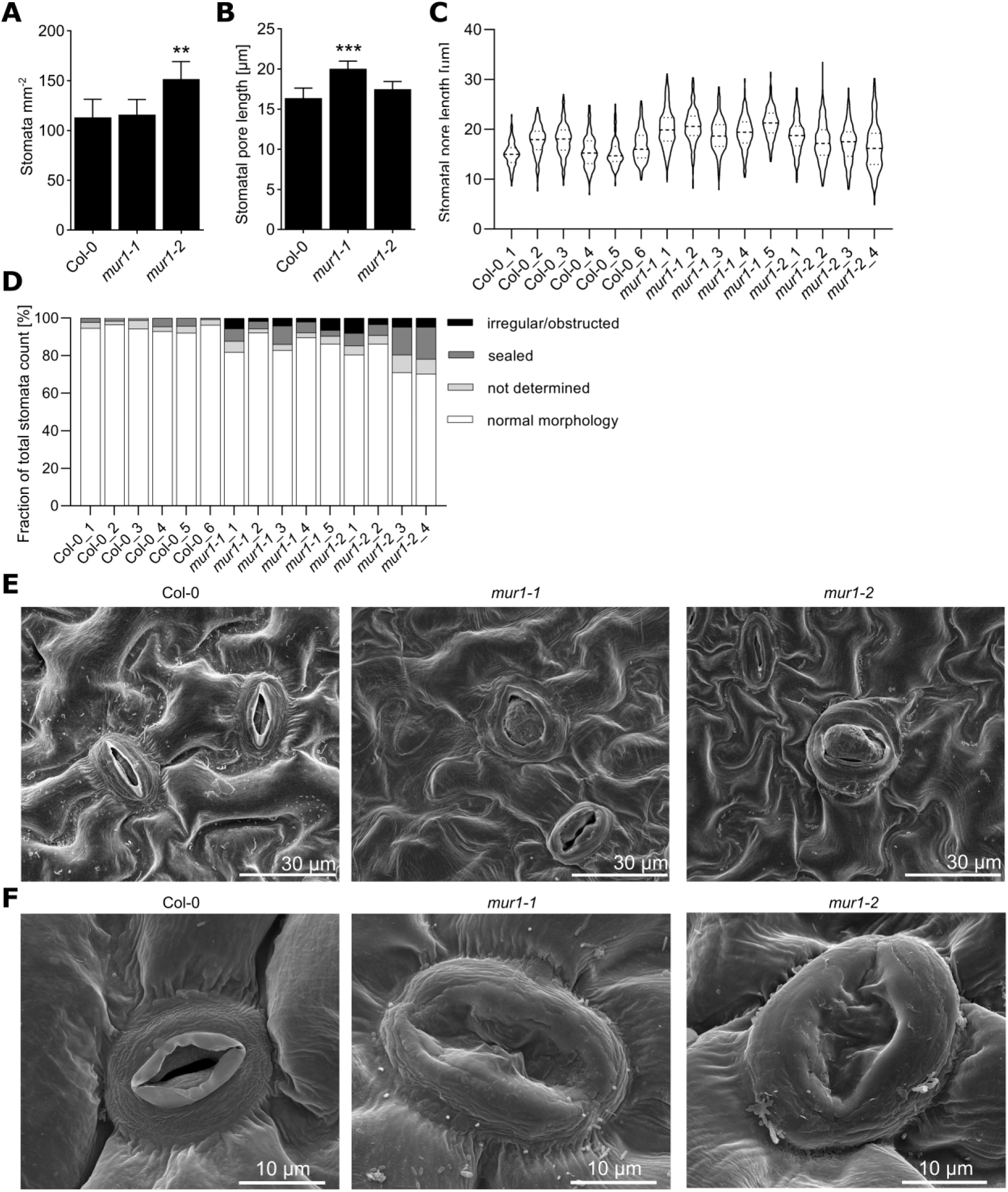
**Scanning electron microscopy-based phenotyping of stomatal morphology in *mur1* mutants.** **(A)** Stomatal density, **(B)** mean stomatal pore length and **(C)** distribution of stomatal pore length in individual plants on abaxial side of 3-week-old cotyledons of Col-0, *mur1-1* and *mur1-2*. **(A, B)** Data bars represent means ± SD (n = 4-6 cotyledons, 1 cotyledon per plant). Asterisks denote statistical differences (** p < 0.01, *** p < 0.001) to Col-0 according to one-way ANOVA followed Dunnett’s post-hoc test. **(D)** Frequency of abnormal stomata in Col-0, *mur1-1* and *mur1-2*. Bar represent frequencies obtained for separate plants. **(A-D)** Experiments were performed three times with similar results. **(E, F)** Representative images of **(E)** irregular/obstructed and **(F)** sealed stomata observed on abaxial side of 3-week-old cotyledons of *mur1* mutants.

**Supplementary Figure 6.**
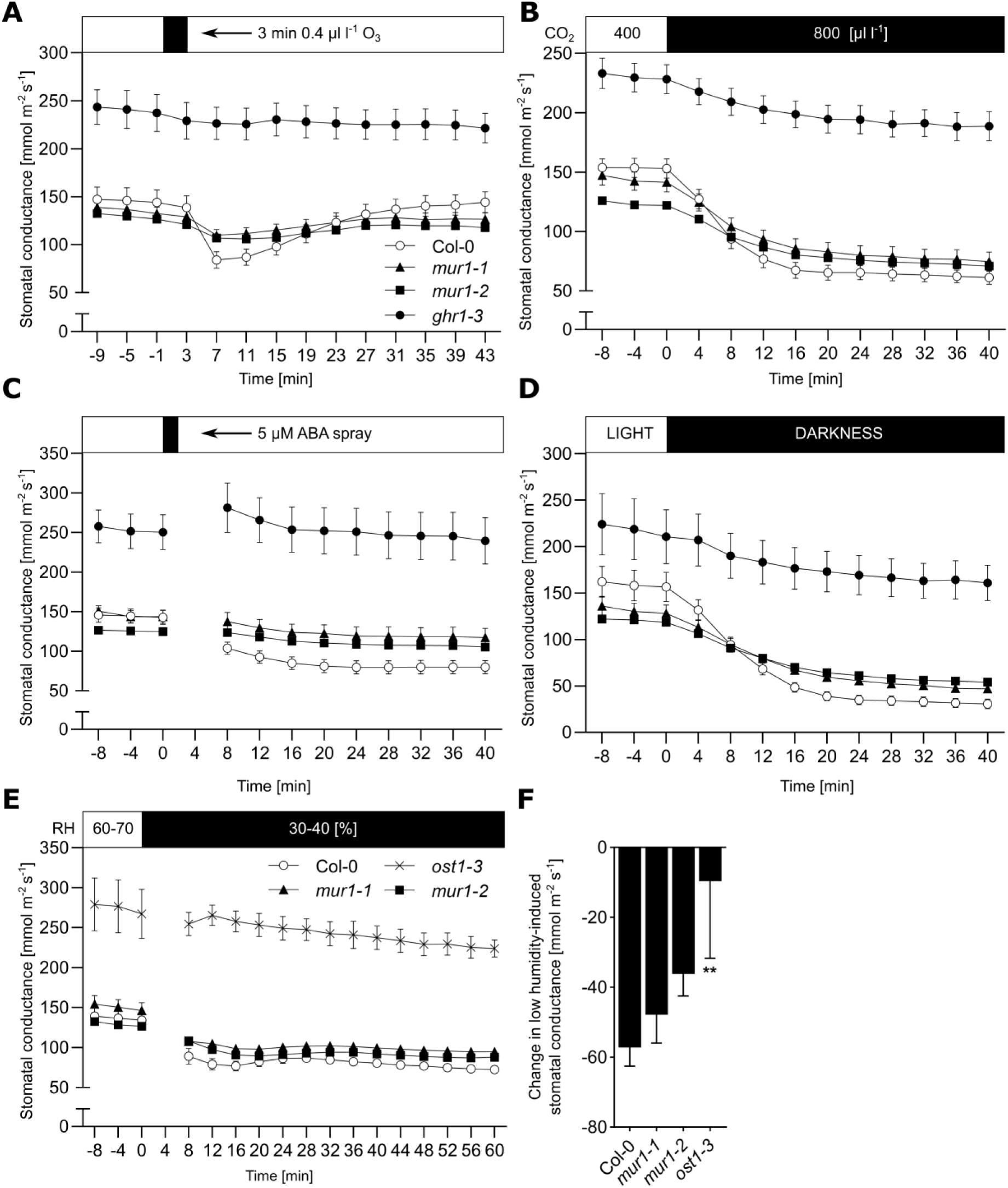
**Stomatal responses of *mur1* mutants to stomata-closing stimuli.** **(A-E)** Whole-rosette stomatal conductance of 3- to 4-week-old *mur1-1*, *mur1-2*, Col-0 and *ghr1-3* **(A-D)** or *ost1-3* **(E)** plants in response to **(A)** O_3_ pulse, **(B)** elevated CO_2_, **(C)** ABA spray, **(D)** darkness, **(E)** drop in relative humidity (VPD increase from 1.01 ± 0.01 kPa to 2.04 ± 0.03 kPa (mean ± SE)). The indicated treatments were applied at t = 0. Data points represent means ± SEM; n = 7-10 **(A)**, 10-12 **(B)**, 9-11 **(C)**, 6-11 **(D),** 3-12 **(E)** plants analyzed in 2 **(A, C, D, E)** or 3 **(B)** independent experiments. **(F)** Changes in low humidity-induced stomatal conductance in Col-0, *mur1-1*, *mur1-2* and *ost1-3*. Values were calculated by subtracting the stomatal conductance at t = 0 from the stomatal conductance at t = 16 min based on data presented in **(E)**. Data bars represent means ± SEM; n = 3-12 plants. Asterisks denote statistical differences to Col-0 (** p < 0.01) according to one-way ANOVA followed by Dunnett’s post-hoc test.

**Supplementary Figure 7.**
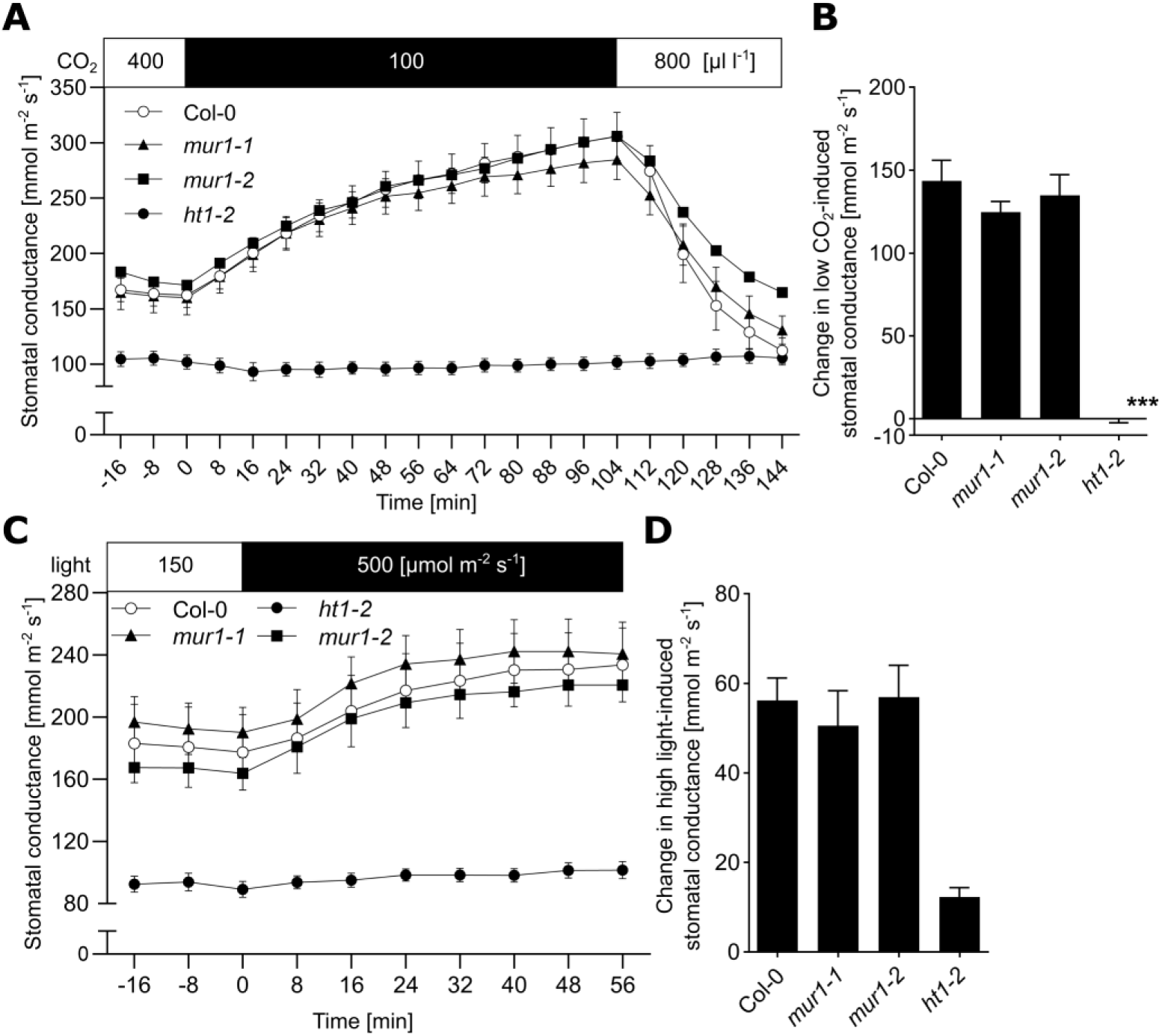
**Response of *mur1* mutants to stomata-opening stimuli.** **(A, C)** Time course of stomatal conductance of *mur1-1*, Col-0 and *ht1-2* plants in response to **(A)** low CO_2_ concentration, **(C)** increase in light intensity. The indicated treatments were applied at t = 0 and whole-rosette stomatal conductance of 3- to 4-week-old plants was recorded. Data points represent means ± SEM; n = 8-10 plants analyzed in 2 independent experiments. **(B)** Change in stomatal conductance 104 min after decrease in CO_2_ concentration, calculated based on the data presented in **(A)**. **(D)** Change in stomatal conductance 56 min after increase of light intensity, calculated based on the data presented in **(C)**. **(B, D)** Values were calculated by subtracting the initial stomatal conductance recorded at t = 0 from the stomatal conductance obtained at **(B)** t = 104 min, **(D)** t = 56 min. Data bars represent means ± SEM; n = 8-10 plants. Asterisks denote statistical differences to Col-0 (*** p < 0.001) according to one-way ANOVA followed by Dunnett’s post-hoc test.

**Supplementary Figure 8.**
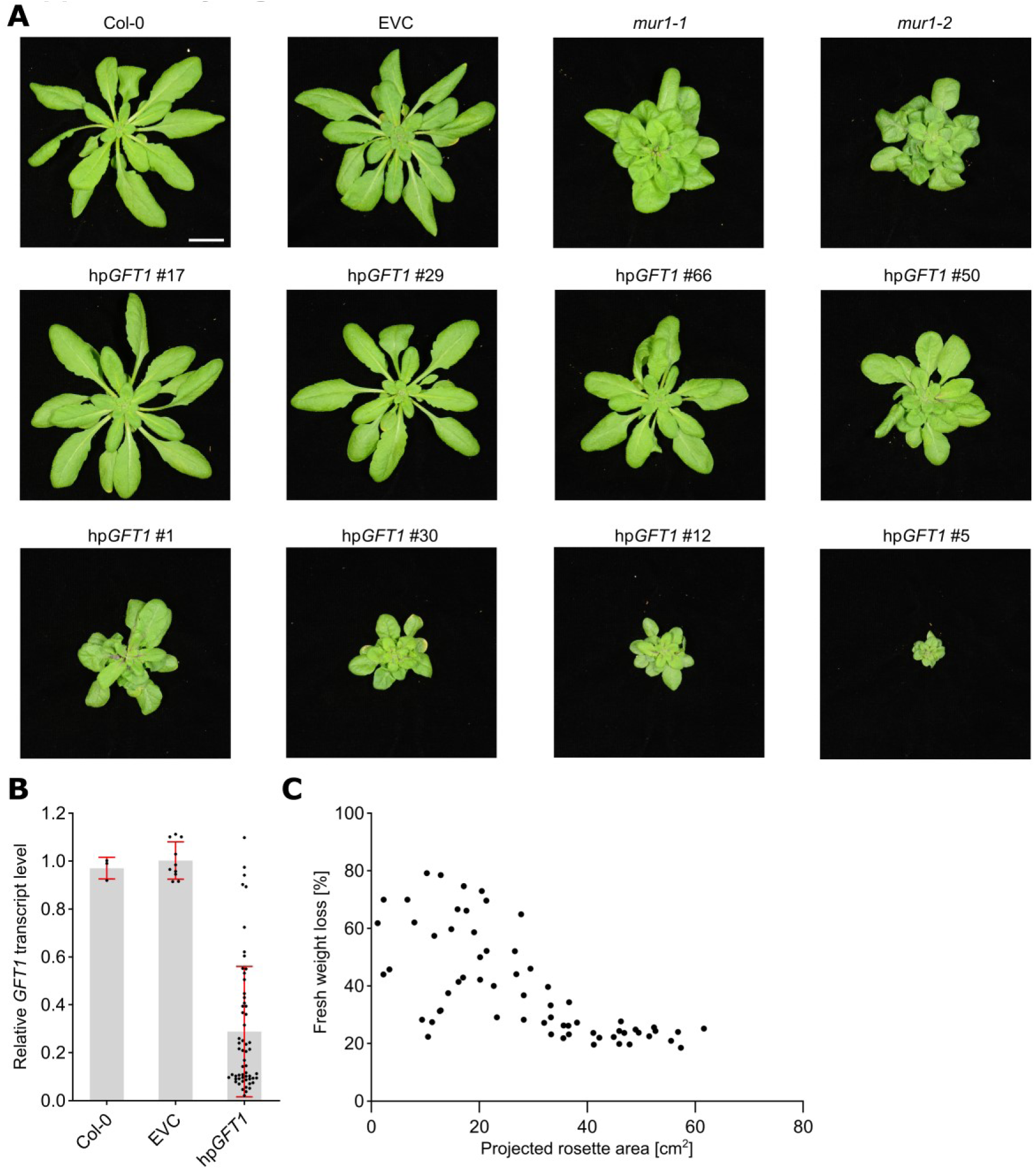
**Analysis of hp*GFT1* T1 plants.** **(A)** Rosette morphology of selected hp*GFT1* T1 plants, empty vector control (EVC), Col-0 and *mur1* mutants. Scale bar = 20 mm. **(B)** Variability in *GFT1* transcript level observed in 58 independent hp*GFT1* T1 plants. Data points represent values obtained for separate plants. Data bars represent means ± SD; Col-0 n = 3 plants, EVC n = 10 plants. **(C)** The relationship between the fresh weight loss in 2h and projected rosette area observed in 64 independent hp*GFT1* T1 plants. Each data point represents values obtained for a single hp*GFT1* T1 plant.

**Supplementary Figure 9.**
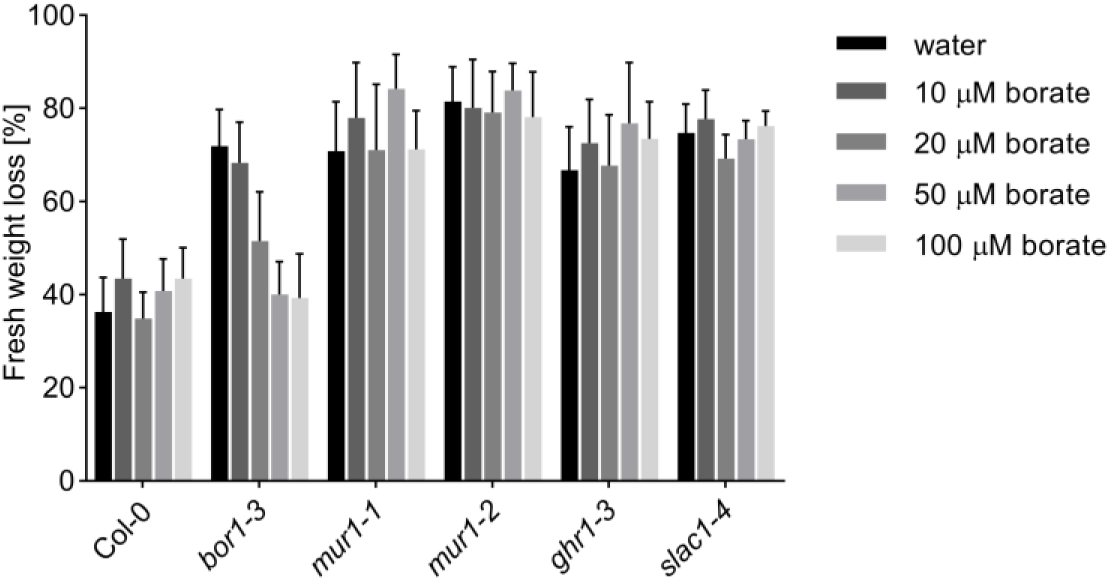
**The influence of soil borate concentration on leaf fresh-weight loss of *bor1-3* mutant.** Data bars represent means ± SD (n = 10-16 plants per genotype per condition). Experiment was performed three times with similar results.

**Supplementary Figure 10.**
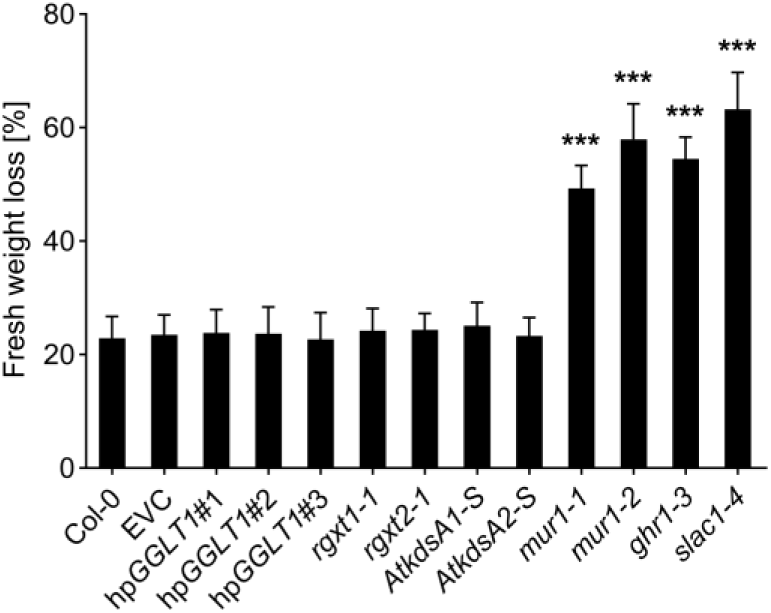
**Water loss of mutants of enzymes/transporters involved in synthesis of RG-II.** Data bars represent means ± SD (n = 13-16 plants). Asterisks denote statistical differences (*** p < 0.001) to respective control lines (Col-0 or EVC-empty vector control) according to one-way ANOVA followed by Sidak’s post-hoc test. Experiment was performed three times with similar results.

**Supplementary Figure 11.**
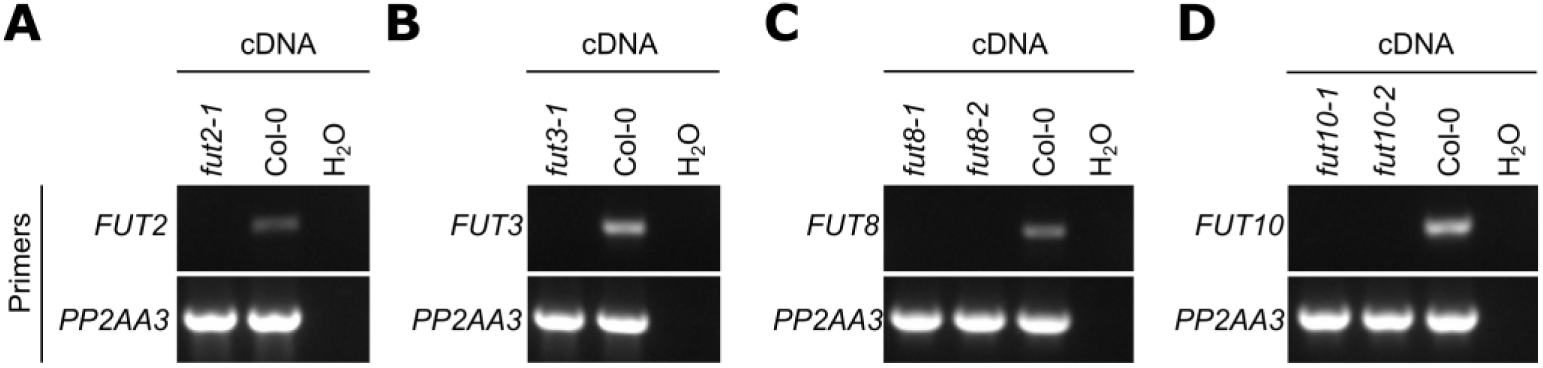
**Characterization of *fut* T-DNA insertion mutants.** **(A-D)** Results of RT-PCR obtained for indicated *fut* mutants with primers specific for **(A)** *FUT2***, (B)** *FUT3*, **(C)** *FUT8* and **(D)** *FUT10*. *PP2AA3* was used as a reference gene and within each panel the same cDNA sample was used for amplification with all primer sets. The experiment was performed with three biological replicates and with similar results. Representative results are shown.

**Supplementary Table 1.**
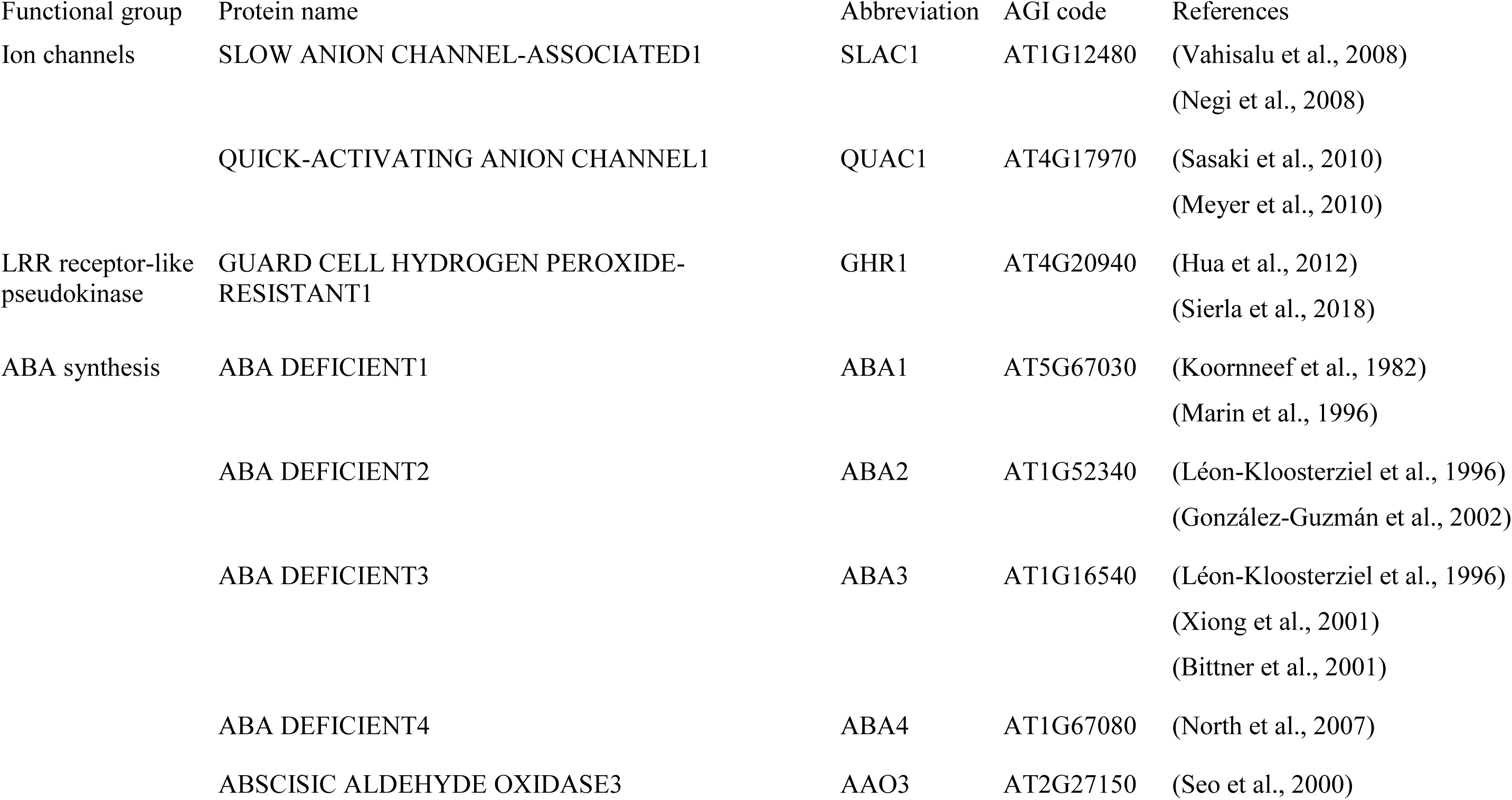
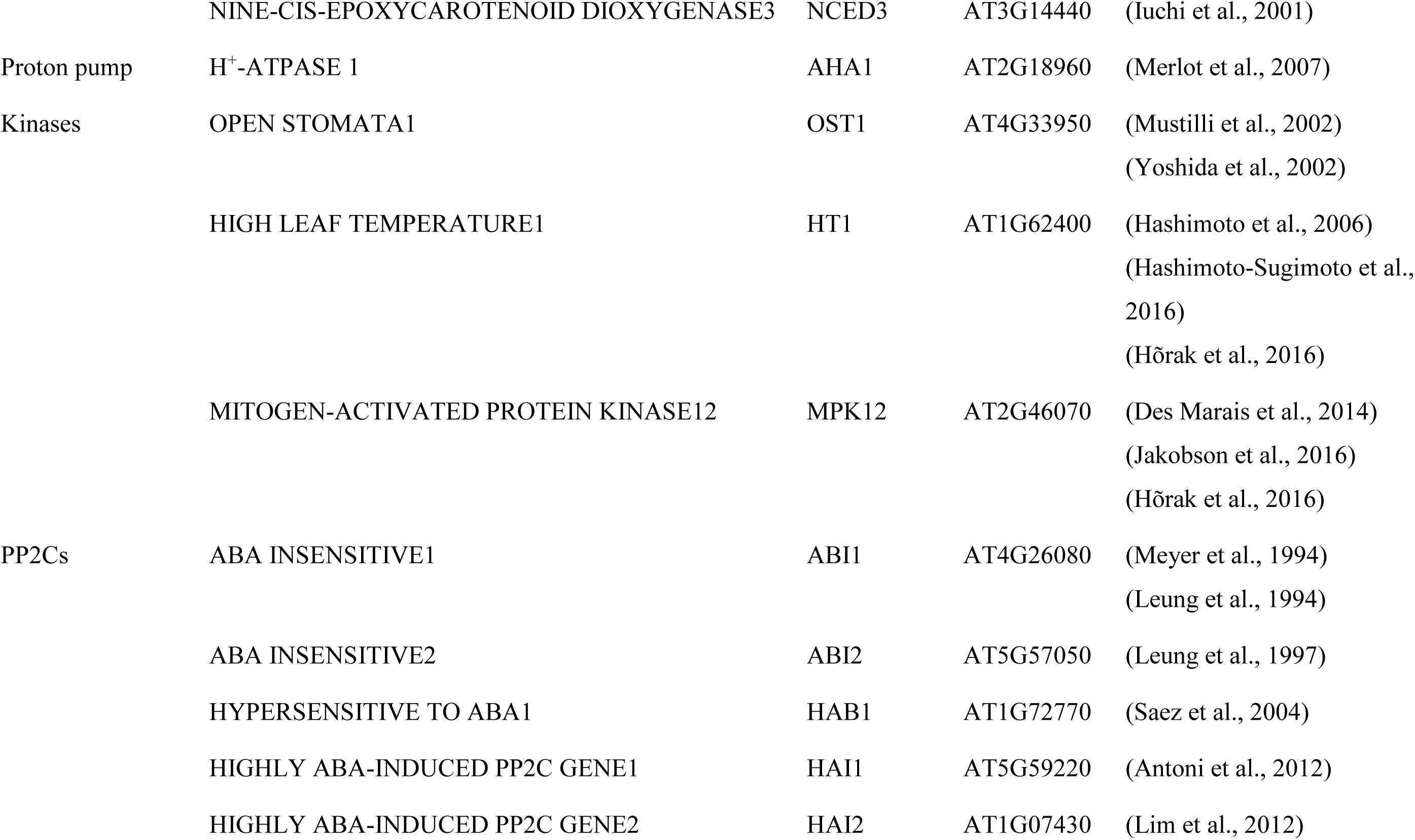
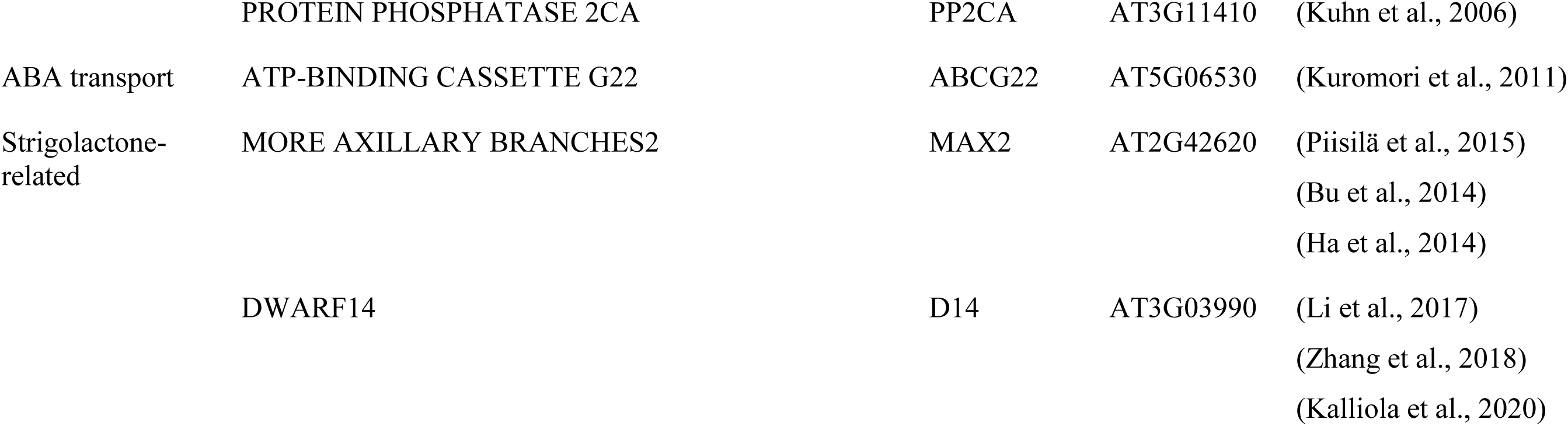
Genes included in candidate gene sequencing.

---

**Supplementary Table 2.** High-frequency single nucleotide polymorphisms identified in T7-9 BC1_F2_ mapping population.

| Chromosome | Mutation | Frequency | AGI code | Reference aa | Mutant aa | Annotation | Mutant lines | Water loss |
| --- | --- | --- | --- | --- | --- | --- | --- | --- |
| 3 | C 19007831 T | 0.95 | At3g51160 | E | K | GDP-D-MANNOSE-4,6-DEHYDRATASE2, GMD2, MUR1, MURUS 1 | *mur1-1* | high |
|  |  |  |  |  |  |  | *mur1-2* | high |
|  |  |  |  |  |  |  | SALK_057153 | high |
| 3 | C 17388552 T | 0.93 | At3g47220 | R | C | ATPLC9, PHOSPHATIDYLINOSITOL-SPECIFIC PHOSPHOLIPASE C9, PLC9 | SALK_025949 | WT-like |
|  |  |  |  |  |  |  | SALK_021982 | WT-like |
| 3 | C 17021471 T | 0.91 | At3g46330 | G | D | MATERNAL EFFECT EMBRYO ARREST39, MEE39 | GK_277G01 | WT-like |
|  |  |  |  |  |  |  | SALK_108641 | WT-like |
|  |  |  |  |  |  |  | SALK_065070 | WT-like |
| 3 | C 18009773 T | 0.91 | At3g48590 | E | K | ATHAP5A, HAP5A, NF-YC1, NUCLEAR FACTOR Y, SUBUNIT C1 | SALK_086334 | WT-like |
| 3 | C 18556988 T | 0.91 | At3g50050 | R | H | Eukaryotic aspartyl protease family protein | SALK_038145 | WT-like |

**Supplementary Table 3.** Genotypes of F3 plants obtained from *ger1^−/−^ger2^+/−^*F2 plants.

| F2 plant | F3 segregation |  |  | n.d.* | n.g.** | Total |
| --- | --- | --- | --- | --- | --- | --- |
|  | *ger1-2*<sup>-/-</sup> *ger2-1*<sup>+/+</sup> | *ger1-2*<sup>-/-</sup> *ger2-1*<sup>+/-</sup> | *ger1-2*<sup>-/-</sup> *ger2-1*<sup>-/-</sup> |  |  |  |
| *ger1-2*<sup>-/-</sup> *ger2-1*<sup>+/-</sup> plant 7/3 | 22 | 23 | - | - | 3 | 48 |
| *ger1-2*<sup>-/-</sup> *ger2-1*<sup>+/-</sup> plant 9/5 | 24 | 22 | - | 1 | 1 | 48 |
|  | *ger1-2*<sup>-/-</sup> *ger2-2*<sup>+/+</sup> | *ger1-2*<sup>-/-</sup> *ger2-2*<sup>+/-</sup> | *ger1-2*<sup>-/-</sup> *ger2-2*<sup>-/-</sup> |  |  |  |
| *ger1-2*<sup>-/-</sup> *ger2-2*<sup>+/-</sup> plant 26/2 | 19 | 20 | - | - | 1 | 40 |
| *ger1-2*<sup>-/-</sup> *ger2-2*<sup>+/-</sup> plant 33/3 | 22 | 18 | - | - | - | 40 |
|  | *ger1-3*<sup>-/-</sup> *ger2-1*<sup>+/+</sup> | *ger1-3*<sup>-/-</sup> *ger2-1*<sup>+/-</sup> | *ger1-3*<sup>-/-</sup> *ger2-1*<sup>-/-</sup> |  |  |  |
| *ger1-3*<sup>-/-</sup> *ger2-1*<sup>+/-</sup> plant 28/2 | 17 | 23 | - | - | - | 40 |
| *ger1-3*<sup>-/-</sup> *ger2-1*<sup>+/-</sup> plant 31/8 | 28 | 11 | - | 1 | - | 40 |
|  | *ger1-3*<sup>-/-</sup> *ger2-2*<sup>+/+</sup> | *ger1-3*<sup>-/-</sup> *ger2-2*<sup>+/-</sup> | *ger1-3*<sup>-/-</sup> *ger2-2*<sup>-/-</sup> |  |  |  |
| *ger1-3*<sup>-/-</sup> *ger2-2*<sup>+/-</sup> plant 37/5 | 19 | 21 | - | - | - | 40 |
| *ger1-3*<sup>-/-</sup> *ger2-2*<sup>+/-</sup> plant 38/6 | 19 | 16 | - | 1 | 4 | 40 |

\* not determined; \*\* not germinated

